# CCR2 is a potential biomarker and therapeutic target in Group 3 medulloblastoma

**DOI:** 10.64898/2026.10.11.757412

**Authors:** Carlos Custódia, Rita Cascão, Eunice Paisana, Daniel Picard, Pedro Ruivo, Tiago Rodrigues, Inês Lourenço, Sofia Nunes, Nan Qin, Luísa Albuquerque, Rafael Roque, José Miguéns, Marc Remke, João T. Barata, Claudia C. Faria

## Abstract

**Background:** Medulloblastoma (MB) is the most common malignant pediatric brain tumor, classified into four molecular subgroups: WNT, SHH, Group 3, and Group 4. Among these, Group 3 MB displays the worst prognosis with the highest rates of dissemination and the worst survival outcomes. While current therapies control the primary tumor, metastatic disease remains largely incurable underscoring the need for novel therapeutic options. The CCL2/CCR2 axis is a well-established metastases driver in systemic tumors through the promotion of angiogenesis and cell migration. Recently, this axis has been implicated in MB pathogenesis and dissemination, particularly in Group 3 tumors. Therefore, we hypothesize that CCR2 represents a therapeutically actionable driver in aggressive MB.

**Methods:** The clinical relevance of CCR2 in MB was assessed using publicly available gene expression data from two independent datasets. CCR2 expression was further evaluated by immunohistochemistry on paired primary and recurrent/metastatic tumor samples to validate these findings. Functional studies using ONS76 and UW228 MB cells with CCR2 overexpression evaluated proliferation, migration and CCR2 downstream signaling. Orthotopic xenograft mouse models assessed CCR2 effect on tumor invasion, dissemination, and survival. An *in-silico* drug screening identified potential CCR2 antagonists that were tested *in vitro* and *in vivo*.

**Results:** Gene expression data analysis across 1094 MB primary samples correlated high *CCR2* expression with dissemination and reduced survival in Group 3 MB patients. In an independent MB patient cohort, CCR2 protein levels were increased in recurrent/metastatic tumors compared with paired primary tumors. *CCR2* overexpression *in vitro* promoted cancer cell proliferation and migration. CCR2-driven MB mouse xenografts, exhibited an aggressive disease phenotype and decreased survival, resembling Group 3 tumors. CCR2 targeting with Cenicriviroc suppressed CCR2-mediated signaling and showed therapeutic efficacy against established MB tumors *in vivo*.

**Conclusions:** *CCR2* identifies aggressive Group 3 MB and represents a promising actionable target for therapy.

## Introduction

Medulloblastoma (MB) is the most common malignant brain tumor in children, accounting for approximately 20% of all pediatric brain tumors and nearly 20% of all childhood central nervous system (CNS) malignancies [1, 2]. MB can be classified into four molecular subgroups, each defined by distinct genetic and clinical features: WNT-activated, SHH-activated, Group 3, and Group 4 [3, 4].

The standard of care treatment for MB includes surgery, followed by craniospinal radiation (for patients older than 3 years) and high-dose chemotherapy [5, 6]. These strategies effectively treat the primary tumor, with a 5-year survival rate ranging from 50% to more than 95% depending on the subgroup [1, 4, 7, 8]. On the other hand, MB metastases fail to respond to current therapies and associate with a dismal patient outcome [9–11]. Among the four MB subgroups, Group 3 carries the poorest prognosis largely due to its high propensity for early dissemination [1, 12], resulting into a 5-year overall survival (OS) below 54% [12]. Since metastatic MB is an incurable disease, there is an urgent need for novel therapeutic targets for treating these patients.

The CCL2/CCR2 pathway has been implicated in MB pathogenesis. CCL2 is a chemokine that preferentially binds to the CCR2 receptor, an axis that is central to immune cell recruitment and is involved in angiogenesis and cell migration [13–15]. CCL2 has been shown to be upregulated in metastatic Group 3 MB patients [16] and was linked to the maintenance of stem-like properties in disseminated MB cells [17]. Moreover, increased CCL2 levels have been observed in the cerebrospinal fluid (CSF) of patients with metastatic Group 3 and Group 4 tumors [18, 19]. Systemically, although high expression of *CCR2* has been linked to decreased survival across various systemic cancers [20–23] and glioblastoma [24], it’s association with MB has not been reported.

The therapeutic targeting of the CCL2/CCR2 axis has been tested in different models of cancer. In a preclinical model of prostate cancer, CCL2 inhibition enhanced treatment response when combined with conventional chemotherapy [25]. CCR2 inhibition showed promising results in cellular models of prostate cancer [26], mouse models of hepatocellular carcinoma [27] and glioma [28]. Furthermore, dual inhibition of CCL2 and CCR2 has been shown to improve therapeutic responses to immune checkpoint inhibitors in preclinical models of lung cancer and glioma [28, 29].

In this study, we hypothesized that targeting CCR2 could provide a therapeutic opportunity for aggressive MB. Preclinical *in vitro* and *in vivo* models of CCR2-driven MB mirrored the aggressive metastatic phenotype encountered in patients. An *in-silico* drug screening identified compounds that successfully targeted CCR2, reducing MB growth and dissemination *in vivo*.

## Materials and methods

### Dataset analysis

*CCL2* and *CCR2* gene expression data from MB patients were obtained from two independent publicly available datasets: Cavalli *et al.* (GSE85218; microarray, n = 763) [30] and Williamson *et al.* (E-MTAB-10767; RNA-seq, n = 331) [31]. Data harmonization was performed in Python 3.13.3 using the pandas, numpy, scipy, mygene, and GEOparse packages. For survival analyses, OS was capped at 120 months. Patients were stratified as *CCL2* or *CCR2* High or Low using a mean expression split computed separately within each cohort. The association between *CCL2* or *CCR2* gene expression and OS were assessed for all MB patients from Group 3 using Kaplan-Meier survival curves with log-rank testing, in each cohort independently. Analyses were repeated within the age strata of 4 to 10 years old and within molecular subtypes. *CCL2* and *CCR2* gene expression levels were compared in Group 3 MB across metastatic status in the strata of 4 to 10 years using pairwise Mann-Whitney post-hoc comparisons. Within Group 3, *CCR2* expression was additionally compared across the three molecular subtypes (alpha, beta, gamma) using Kruskal-Wallis test. All statistical analyses were performed in R 4.5.2 using the survival and broom packages. Visualizations were produced using the ggplot2, ggbeeswarm, ggrepel, patchwork, survminer, and svglite packages.

### Histopathological analysis of MB patients

The use of human samples in this study was performed in accordance with the Unidade Local de Saúde Santa Maria Ethics Board (Refa. N° 367/18 and Refa. N° 346/20). Eleven paired formalin-fixed paraffin embedded (FFPE) samples of primary tumors and metastases or recurrent tumors from MB patients were kindly provided by the Neuropathology Laboratory at Hospital de Santa Maria, Unidade Local de Saúde Santa Maria (HSM-ULSSM). Tissue samples were serially sectioned at a thickness of 3μm using a microtome, mounted on microscope slides and stained with hematoxylin and eosin (H&E) for morphological examination. Tissue sections were also used for immunohistochemical (IHC) staining following conventional protocols, using the CCR2 antibody (1:100; Abcam, ab231039). Briefly, antigen retrieval slides were treated in a PT Link module (DAKO, USA) at low-Ph, followed by incubation with the primary antibody. EnVision Link horseradish peroxidase/DAB visualization system (Dako, USA) was used, and sections were then counterstained with hematoxylin and mounted. Evaluation of IHC was performed by two independent researchers using semi-quantitative scores of staining intensity (low or high staining intensity) and frequency (low: 0%-49% or high: 50%-100%). The results were expressed as a combination of both intensity and frequency divided into four levels: level I (low frequency/low intensity), level II (low frequency/high intensity), level III (high frequency/low intensity), and level IV (high frequency/high intensity); and were validated by a neuropathologist. Representative photomicrographs were acquired using a Leica DM2500 microscope coupled to a Leica MC170 HD microscope camera (Leica Microsystems, Germany).

### Cell Lines and Culture Conditions

Human MB cell lines (ONS76 and UW228), previously engineered to stably express GFP and Luciferase (GFP-Luc), were kindly provided by Dr. James Rutka, at The Hospital for Sick Children, Toronto, Canada. HEK-293T cells were purchased from ATCC. All cell lines were cultured in DMEM 1x (Gibco, 41966-029), supplemented with 10% FBS (BioWest, S1810-500), 2mM L-glutamine (Gibco, 25030-024), 1% HEPES (Gibco, 15630-080), and 1% Pen-Strep (Gibco, 15070-063) until passage 30. Cells were divided when 70-90% confluency was reached. To passage the cells, the supernatant was discarded, and cells were washed with PBS 1x. Cells were then incubated for three minutes with Trypsin-EDTA 0.05% (Gibco, 15400-054) to induce cell detachment from the flask. The resulting cell suspension was diluted with DMEM and divided into new culture flasks.

### Stable Overexpression of *CCR2*

ONS76 and UW228 cell lines (GFP-Luc) were genetically engineered to stably overexpress (OE) *CCR2* variant B through lentiviral vector gene delivery. A lentiviral plasmid encoding the open reading frame of *CCR2* variant B (Origene, RC225538L3) was used in this process. An empty vector plasmid (Origene, PS100092) was used as the control for transduction. For viral production, HEK-293T cells were seeded at a density of 2×10⁶ cells per plate onto 10cm² culture dishes 24 hours prior to transfection with the plasmid cocktail for viral production. Briefly, 24 hours after seeding, cells were exposed to a combination of the lentiviral vector, envelope, and capsid plasmids, together with lipofectamine 2000 (Invitrogen, 11668030) to enhance transfection efficiency. HEK-293T supernatant, containing viral particles, was collected at 48- and 72-hours post-transfection and filtered through a 0.45 μm filter prior to its use in infection. ONS76 and UW228 MB cells (GFP-Luc) were infected with the produced virus by incubation with the HEK-293T supernatant in the presence of 8 μg/ml polybrene (Sigma-Aldrich, TR-1003). Selection of stably transduced cells was achieved through selection with puromycin (InVivoGen, Ant-pr-1), with a concentration of 1.5 μg/ml for ONS76 cells and 1.0 μg/ml for UW228 cells. Additional purification of *CCR2* overexpressing cells was performed through flow cytometry sorting using BD FACSAria III cell sorter. Confirmation of *CCR2* overexpression was performed by flow cytometry using a phycoerythrin (PE)-conjugated anti-CD192 (CCR2) antibody (1:100; Miltenyi Biotec, 130-118-477; RRID:AB_2751523). The resulting *CCR2*-overexpressing cell lines were named ONS76 OE-CCR2 and UW228 OE-CCR2. Control cell lines transduced with empty vector were named ONS76 control and UW228 control.

### Surface CCR2 quantification by flow cytometry

CCR2 levels were assessed by flow cytometry. Briefly, ONS76 OE-CCR2 and UW228 OE-CCR2 cells were seeded onto a 6-well plate at 3×10^5^ cells per well. ONS-76 control and UW288 control cells were used to assess baseline expression levels. Twenty-four hours after seeding cells were starved in DMEM media with 0.1% FBS overnight. Adherent ONS76 and UW228 cells were detached using a Trypsin-EDTA 0.05% (Gibco, 15400-054), washed with PBS 1x and centrifuged at 300g for 5 minutes. Cells were then resuspended in a solution containing a phycoerythrin (PE)-conjugated anti-CCR2 antibody (Miltenyi Biotech, 130-118-477) diluted 1:100 in PBS 1x with 2% FBS and incubated for 20 min at room temperature in the dark. After incubation, cells were washed and resuspended in PBS 1x with 2% FBS. Percentage of positive CCR2 cells and the mean fluorescence intensity (MFI) of CCR2 was measured by flow cytometry using a BD LSRFortessa Cell Analyzer.

### Cell proliferation

The CellTiter 96 Aqueous One Solution Reagent (MTS; Promega, G3580) was used to assess proliferation, as defined by the manufacturer’s protocol. Briefly, modulated MB cells (both control and OE-CCR2) were seeded in 96-well plates, at 500 cells per well in appropriate culture media. In each time-point, cells were incubated for 2 hours with MTS (Promega, G3580) and the absorbance was measured at 490 nm using a microplate reader Tecan Infinite M200 (Tecan, Austria). Cell proliferation readings were acquired at 0, 24, 48, 72, and 96 hours. The fold change in proliferation was determined by normalizing the results to 0 hours. Three independent experiments were performed, with three technical repetitions each. In addition, to assess the stimulatory effect of exogenous CCL2 on cell proliferation 50 ng/mL of recombinant human (rh)CCL2 (Abcam, ab270081) was added to the culture media at 0 hours and the experiment was performed as described above.

### Cell migration

Migration assays were performed using Boyden chambers (8 μm pore size; Corning, 80209). Fifty thousand ONS76 OE-CCR2 or UW-228 OE-CCR2 cells were seeded in the top chamber in starvation media (DMEM+0.1%FBS). The bottom chamber was loaded with starvation media supplemented with 50ng/mL of rhCCL2. Cells were allowed to migrate for 16 hours (ONS76 OE-CCR2) or 36 hours (UW-228 OE-CCR2) at 37°C and 5%CO_2_. Then, non-migrating cells were removed from the top side of the chamber and cells that migrated through the porous membrane (on the bottom side) were stained with DAPI (Thermo Fischer Scientific, D1306) and photographed with an EVOS M5000 (Invitrogen) microscope. The number of cells was determined by the analysis of 5 random fields (10x magnification) using ImageJ. Three independent experiments were performed, with three technical replicates each.

### Immunoblotting

ONS76 OE-CCR2 MB cells were seeded in 6-well plates at 3×10⁵ cells per well. Twenty-four hours after seeding, cells were starved in DMEM media with 0.1% FBS overnight. Then, to assess the kinetics of CCR2 signaling upon CCL2 stimulation, cells were incubated with 50ng/mL of rhCCL2 (Abcam, ab270081) for 1, 5, 15, 30 and 60 minutes prior to cell lysate collection. Cell lysates were prepared by directly adding to plated cells a lysis buffer containing 50mM Trizma-Base (Sigma-Aldrich, 93350), 150mM NaCl (Sigma-Aldrich, S9888), 5mM EDTA (Sigma-Aldrich, 03609), 1% (v/v) IGEPAL® CA-630 (Sigma-Aldrich, I8896), supplemented with 1x PhosSTOP™ phosphatase inhibitors (Roche, 4906845001) and 1x cOmplete™ protease inhibitor cocktail (Roche, 11836153001). After centrifugation at 10,000g for 15 minutes at 4°C, the supernatant was harvested. Total protein concentration was determined using the Bradford protein assay (Protein Assay Dye Reagent Concentrate; BioRad, 5000006). Equal amounts of protein were separated by SDS-PAGE on 12% Acrylamide/Bis-acrylamide 29:1 gels, 120V, 90min, and transferred onto nitrocellulose membranes (BioRad, 1620112) at 400mA, 90min, 4°C. Membranes were then blocked with 3% skim milk in Tris Buffer Saline with 0.1% Tween-20 (TBS-T) (Sigma-Aldrich, P1379) for 1 hour at room temperature. Membranes were incubated with the primary antibodies overnight at 4°C and then incubated with the appropriate secondary antibodies (anti-rabbit IgG, HRP Conjugate, Promega, W4011; anti-mouse IgG, HRP Conjugate, Promega, W4021) dilution at 1:5000 for 1 hour at room temperature. After incubation, membranes were washed in TBS-T and incubated with Pierce™ ECL Western Blotting Substrate (Thermo-Fisher Scientific, 32106). Exposed film (Agfa, CP-BU) was developed using an Agfa Curix 60 automated film processor. Antibodies against p-S473-AKT (1:1000; Cell Signaling Technologies, 4060), AKT (1:1000; Cell Signaling Technologies, 9272), p-p44/42 MAPK (pERK; 1:1000; Cell Signaling Technologies, 9101), p44/42 MAPK (ERK; 1:1000; Cell Signaling Technologies, 9107), and beta-actin (1:1000; Abcam, ab8226) were used. Beta-actin was used as loading control, referred simply as actin throughout the manuscript. Two independent experiments were performed. Densitometry analysis was performed using ImageJ software. Each band was analyzed with a constant frame.

### Orthotopic MB mice models

All mice studies were approved by the institutional animal welfare body (ORBEA-GIMM) and conducted in accordance with Directive 2010/63/EU (transposed to Portuguese legislation through Decreto-Lei N°. 113/2013, of August 7^th^) to ensure that the use of animals complies with all applicable legislation and follows the 3R’s principle. This study was licensed by the Portuguese competent authority (Direcção Geral Animal e Veterinária, DGAV; DGAV Project License Numbers: 012028 \2016, and 017724 \2021). All animals were kept in specific pathogen-free (SPF) conditions, randomly housed per groups under standard laboratory conditions (at 20-22°C under 10-hour light/14-hour dark), and given free access to food (RM3, SDS Diets, Witham, UK) and water (Ultrapure).

Six to 12 weeks of age NOD.Cg-Prkdcscid Il2rgtm1Wjl/SzJ (NSG) mice were purchased from Charles River Laboratories or obtained from a NSG colony established in-house. All surgical procedures were performed under volatile anesthesia (Isoflurane, B. Braun Medical) and treatment with 100µL of 15µg/mL buprenorphine (Bupaq, Plurivet). ONS76 OE-CCR2 (100.000 cells in 3μL of PBS) or UW-228 OE-CCR2 (250.000 cells in 3μL of PBS) cells were injected intracranially, using the lambda as a reference point (2mm lateral to the right, - 1mm anterior, and 2.5mm ventral). After surgery, mice were allowed to recover on a heating pad and monitored for any signals of distress. To assess survival, mice were allowed to live until reaching at least one humane endpoint. Time-point experiments were also performed up to 25 days after intracranial injection using the ONS76 OE-CCR2 cells. Eight to 15 female NSG mice were used per group in all experiments. Mice were monitored for body weight, discomfort, and distress every other day. Humane endpoints were established including 20% body weight loss, head-tilt, neurological disorders (such as paralysis and stereotypic behavior), and tumor outgrowth palpable in the skull. Once they reached the humane endpoint, animals were euthanized by intraperitoneal injection of 140mg/kg sodium pentobarbital (Euthasol, Dechra). During necropsy, whole CNS (brain and spinal cord) was collected for histopathological analysis.

### Histopathological analysis of mouse samples

Mice tissue samples were obtained at necropsy and fixed immediately in 10% neutral buffered formalin solution, dehydrated and embedded in paraffin. Samples containing bone were decalcified in 10% formic acid. For morphological examination, serial 3 µm sections were stained with H&E and representative photomicrographs were taken using Leica DM2500 brightfield microscope coupled to a Leica MC170 HD microscope camera (Leica Microsystems, Germany). To better understand the dissemination and invasion patterns of modulated MB cells in the CNS, we analyzed H&E-stained brain and spinal cord tissue sections using semi-quantitative scoring systems. Dissemination score: 0 – Negative (no dissemination); 1 – Low (minimal to mild); 2 – High (marked). Invasion score: 0 – Negative (no tumor invasion); 1 – Low (minimal to mild); 2 – High (marked). Histopathological analysis was performed by two independent researchers and validated by a specialized neuropathologist.

### *In-silico* drug screen

To identify potential inhibitors specific of CCR2, the Drug–Target Relationship Predictor (DEcRyPT) was used [32, 33]. This tool employs a random forest (RF) regressor and type scaled CATS2 topological pharmacophore descriptors (MOE implementation). DEcRyPT was built using manually and automatically ChEMBL26 curated data to solely include relevant bioactivity information for model assembly, as previously described. This tool predicted more than 10,000 compounds, of which we selected the top 50 with higher affinity rank for CCR2. The top 50 compounds were further narrowed down based on available published data and specific biochemical and pharmacological properties, such as drug class. The top 3 selected candidate compounds were Cenicriviroc (referred as CVC; a dual CCR2/CCR5 inhibitor; Selleck Chemicals, S8512), Marizomib (referred as MAR; a proteasome inhibitor; Merck, SML1916) and Romidepsin (referred as ROM; an HDAC inhibitor; Selleck Chemicals, S3020).

### *In vitro* drug testing

To determine the effect of CVC, MAR and ROM treatment on MB cells *in vitro*, we assessed the CCR2 levels at cell surface, CCR2 downstream signaling, and cell viability, proliferation and migration. These *in vitro* assays were performed using the following drug concentrations: 5µM, 10µM or 50µM of CVC; 50nM or 100nM of MAR; and 2.5nM, 5nM or 10nM of ROM. These concentrations were selected based on previous published studies [34–36]. Stock solutions for CVC, MAR and ROM were prepared in dimethyl sulfoxide (DMSO).

To assess the surface levels of CCR2, cells were incubated with the above-mentioned drug concentrations for 24, 48, 72 and 96 hours (CVC) or for 8, 14, 24 and 48 hours (MAR and ROM). After incubation cells were stained with CCR2-PE and analyzed by flow cytometry, as described above.

To assess the CCR2 signaling, ONS76 OE-CCR2 cells were incubated for 2 hours with 5µM or 10µM of CVC, 50nM of MAR, and 10nM ROM prior to rhCCL2 (50ng/mL) stimulation. Cell lysates and immunoblotting were performed as described above.

To measure viability, cells were incubated with the above-mentioned drug concentrations for 24, 48, 72 and 96 hours (CVC) or for 8, 14, 24 and 48 hours (MAR and ROM). After incubation cells were stained with a permeant fixable LIVE/DEAD™ Fixable Far Red Dead Cell Stain (Life Technologies, Thermo Fisher Scientific, Cat#L10120) for 30min at RT. Percentages of live cells were measured by flow cytometry.

To assess proliferation, cells were incubated with the above-mentioned drug concentrations, in the presence or absence of 50ng/mL of CCL2, and an MTS assay was performed until 96 hours, as described above. In the case of CVC, proliferation was determined by manual cell counts using the trypan blue exclusion method and a Neubauer chamber. ONS76 OE-CCR2 were incubated with 10µM or 50µM CVC and UW-228 OE-CCR2 were incubated with 10µM, 25µM or 50µM CVC.

To evaluate migration, cells were incubated with 50μM of CVC and the Boyden chamber migration assays were performed, as described above.

Three independent experiments were performed in all assays, with three technical replicates each. DMSO was used as a negative control for comparison.

Initial *in vitro* testing of the three candidate compounds was performed in ONS76 OE-CCR2 cells. Based on our results, CVC was identified as the most promising and subsequently, the only compound evaluated in UW-228 OE-CCR2 cells.

### *In vivo* drug testing

Drug efficacy was evaluated *in vivo* using orthotopic MB mice models. NSG mice were injected with ONS76 OE-CCR2 cells, as described above, and treated with 30mg/kg/day of CVC (dose based on previous studies[37, 38]). CVC was freshly dissolved in 10% hydroxypropyl-β-cyclodextrin (Sigma-Aldrich, Cat#332593), 5% kolliphor (Sigma-Aldrich, Cat#07076), and 85% sterile water, immediately before administration by intraperitoneal injection.

Animals were randomized into 2 groups: CVC-treated (N=10) and control (vehicle; N=15). Treatment started five days after intracranial injection of cells and mice received three cycles of therapy (each cycle with 5 days of treatment followed by a wash-out period of 2 days) until day 24 post-injection.

Mice were monitored every other day, and body weight variations were recorded throughout treatment. Mice were euthanized 24h after the last treatment administration (at day 24), and CNS was collected for histopathological analysis, as described above.

### Mice *in vivo* imaging

Mice were injected subcutaneously with XenoLight D-Luciferin, Potassium Salt (Perkin Elmer, PELS122799), 10 min before image acquisition. Animals were imaged under anesthesia (100 mg/kg ketamine + 10 mg/kg xylazine cocktail in saline, delivered intraperitoneally at 0.1 mL per 10g body weight). Images were acquired from the dorsal region to enhance the capture of signals derived from the CNS (brain and spinal cord). *In vivo* bioluminescent images were acquired with 5 minutes of exposure, immediately before mice euthanasia and necropsy using an IVIS Lumina System (Perkin Elmer). Bioluminescence signal intensity quantification was performed using the Living Image® software (PerkinElmer/Revvity) by defining a region of interest (ROI) encompassing the animal CNS.

### Statistical analysis

Normality distribution was assessed by D’Agostino and Pearson test and statistical differences were determined using non-parametric Mann–Whitney tests. Survival curves were analyzed using log-rank tests (Mantel-Cox). CCR2 IHC semi-quantitative scores in patient samples were compared between primary and metastatic/recurrent tissues using a Chi-Squared test. Histological scores from *in vivo* experiments were compared between conditions using ordinal logistic regression (proportional odds model, MASS package v7.3-65). These analyses were performed in R (version 4.6.0). All other statistical analyses were performed using the GraphPad Prism v6.0 (GraphPad, California, USA, RRID:SCR_002798). Bar graphs represent median and interquartile range. Differences were considered statistically significant for p<0.05. Sample size calculations were performed for *in vivo* experiments using the G*Power 3.1 software (Heinrich-Heine University of Dusseldorf).

## RESULTS

### *CCR2* identifies aggressive Group 3 MB and associates with worse patient survival

To determine the clinical significance of *CCR2* and *CCL2* gene expression levels in Group 3 MB patients, we analyzed 1094 MB primary samples in publicly available datasets from Cavalli *et al.* (GSE85218; microarray) [30] and Williamson *et al.* (E-MTAB-10767; RNA-seq) [31]. Higher expression of CCR2 was associated with significantly decreased patient survival in both independent cohorts (**Figure 1A**, Cavalli cohort, p=0.025; **Figure 1B**, Williamson cohort, p=0.024). Refinement of the analysis according to the pediatric age range from 4 to 10 years old [3, 10], identified an association with a worse OS in both independent cohorts (**Figure 1C** Cavalli cohort, p=0.007, **Figure 1D** Williamson cohort, p=0.049). Also, there was no association between *CCL2* expression and patient outcome in both datasets (**Figure S1A-D**). We observed that *CCR2* and *CCL2* displayed significantly increased expression in metastatic patients (M1) when compared to the non-metastatic group (M0) (*CCR2*, p=0.0483 **Figure 1E**; *CCL2*, p= 0.0361, **Figure S1E**). Since *CCL2* expression was not associated with the patients’ survival outcome, we proceeded to a deeper analysis focused on *CCR2*. We did not observe differences in *CCR2* expression across the Group 3 MB subtypes, (**Figure S2A**) and there was no association between high *CCR2* expression for subtypes alpha (α, p=0.912, **Figure S2B**) or beta (β, p=0.054, **Figure S2C**). However, in MB subtype gamma (γ) high *CCR2* expression associates with decreased patient survival (p=0.038, **Figure S2D**).

**Figure 1.**
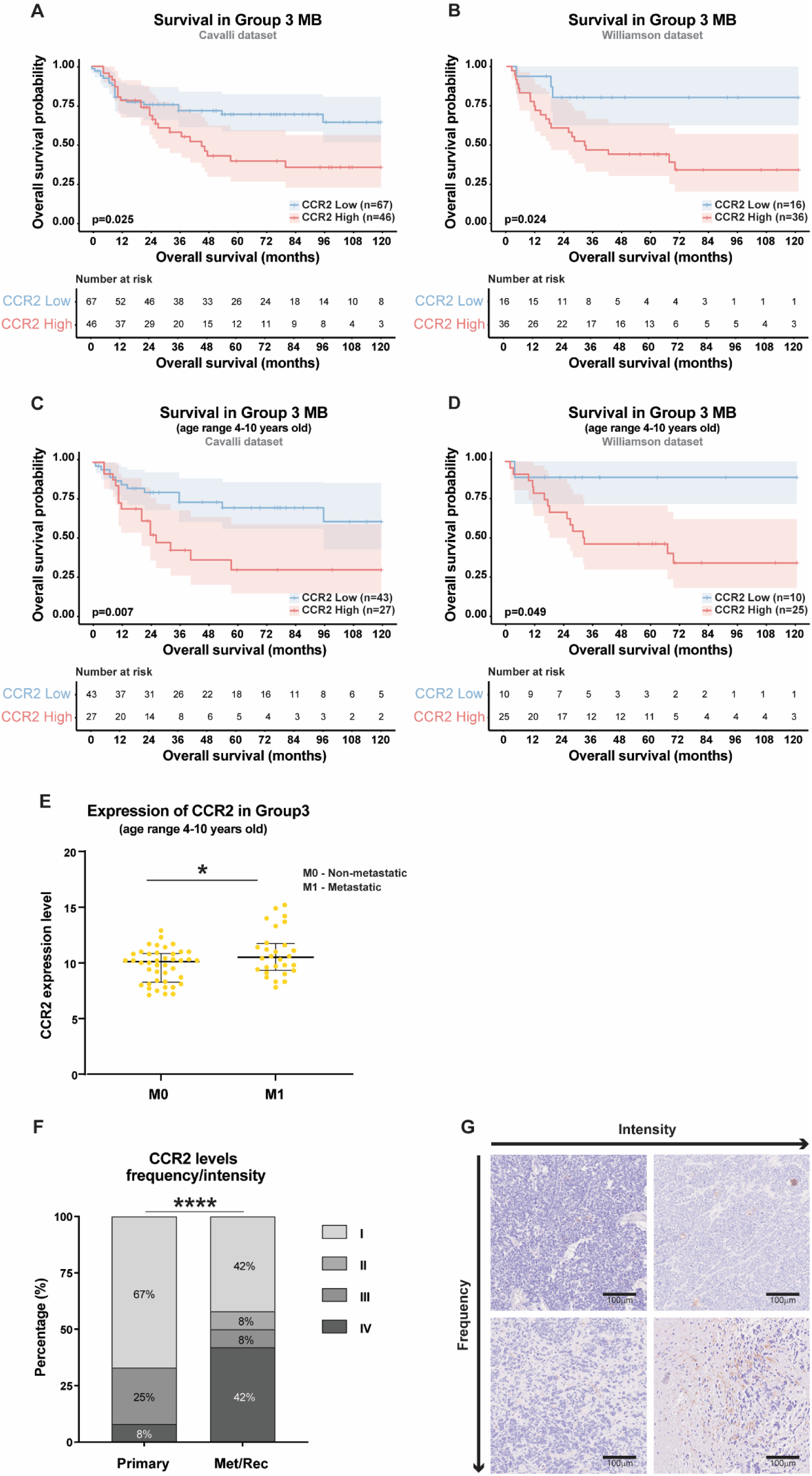
*CCR2* expression is associated with Group 3 MB worse patient outcome. Kaplan-Meier analysis of overall survival (OS) stratified by CCR2 expression levels across Group 3 MB at all ages in the Cavalli **(A)** and Williamson **(B)** datasets. Kaplan-Meier analysis of OS, stratified by CCR2 expression levels, across Group 3 MB int the 4 - 10 years old age range in the Cavalli **(C)** and Williamson **(D)** datasets. Shaded areas represent 95% confidence intervals and statistical significance was determined using the two-tailed log-rank test. (**E**) In the Cavalli dataset *CCR2* is highly expressed in metastatic (M1) cases compared to non-metastatic (M0) cases (p=0.0483). Statistical significance was determined using the non-parametric Mann-Whitney test. (**F**) Semi-quantitative analysis of CCR2 protein expression in a cohort of 11 paired primary and metastatic/recurrent MB samples (p<0.0001). Statistical significance was determined using the Chi-Square tests. (**G**) Representative IHC images illustrating the four-tier scoring system used to evaluate staining: level I (low frequency/low intensity), level II (low frequency/high intensity), level III (high frequency/low intensity), and level IV (high frequency/high intensity). Scale bars represent 100 μm. Significance levels: *p<0.05, **p<0.01; ****p<0.0001; ns, not significant.

*CCR2* findings were validated using an independent cohort of 11 MB patients, submitted to surgical resection at Hospital de Santa Maria (HSM-ULSSM), with paired samples from primary and recurrent/metastatic tumors (**Table S1**). Patients in this cohort had a median age of 12 years (6 - 24 years old), with a higher frequency in males than females (73% and 27%, respectively). IHC analysis revealed a significant increase in CCR2 levels in metastatic and recurrent tumors when compared to the paired primary MB tumor (**Figure 1F and 1G**). These findings suggest that *CCR2* may have prognostic value in Group 3 MB patients.

### CCR2 activation increases MB cell proliferation and migration *in vitro*

The functional role of CCR2 activation in MB was evaluated *in vitro* using MB cell lines engineered to stably overexpress *CCR2* (ONS76 OE-CCR2, representative of Group 3/4, and UW228 OE-CCR2 cells, representative of SHH). Confirmation of *CCR2* expression was assessed by flow cytometry (**Figure S3A-D**).

To assess the functional role of CCL2/CCR2 pathway activation in MB, we performed proliferation and migration assays. Overexpression alone did not influence ONS76 OE-CCR2 proliferation. However, when stimulated with 50ng/mL of CCL2, these cells significantly increased proliferation (24 hours, *p=0.0007*; 48 hours, *p=0.0007*; 72 hours, *p=0.0043*; 96 hours, *p=0.0012;* **Figure 2A**). Also, stimulation with CCL2 increased migration in these cells (*p<0.0001;* **Figure 2B**). On the other hand, overexpression of CCR2 in UW228 cells increased proliferation (48 hours, *p=0.0087*; 72 hours, *p=0.0043*; 96 hours, *p=0.0022*; **Figure S3E**), an effect that was further enhanced upon stimulation with 50ng/mL of CCL2 (72 hours, *p=0.026*; 96 hours, *p=0.0043*; **Figure S3E**). However, CCL2 stimulation did not promote migration in these cells (**Figure S3F**). The downstream signaling of CCL2/CCR2 axis was analyzed by immunoblotting. We assessed the kinetics of CCL2 stimulation in ONS76 OE-CCR2 cells by quantifying activation of ERK and AKT. We observed that ERK phosphorylation increased (**Figure 2C and 2D**), but not AKT phosphorylation (**Figure 2C and 2E**), following CCL2 stimulation, peaking at 1 min and returning to basal levels after 15 min.

**Figure 2.**
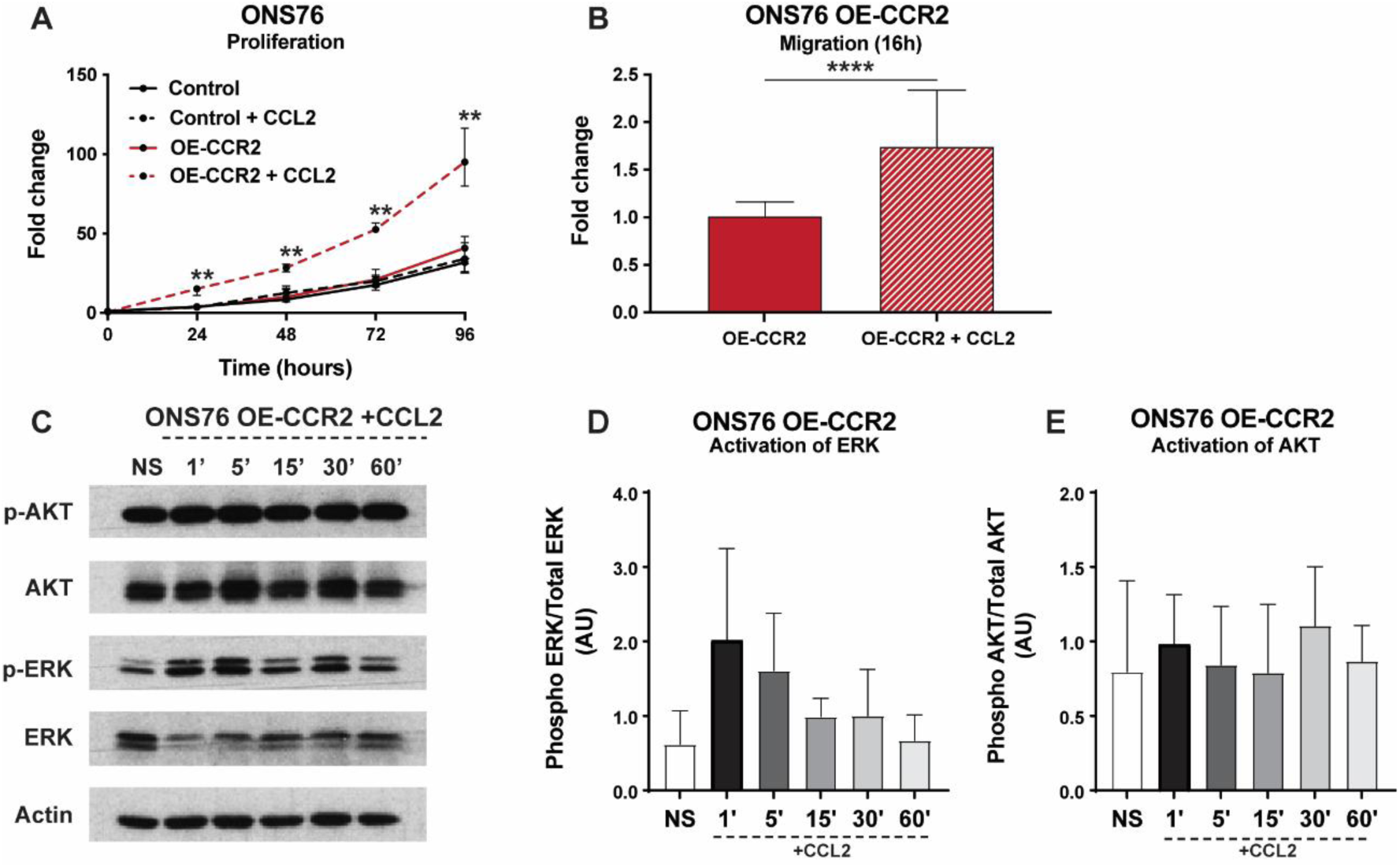
MB cells overexpressing CCR2 respond to CCL2 stimulation, increasing proliferation, migration and ERK activation. (**A**) Proliferation kinetics (fold change to 0h) of ONS76 cells measured by MTS assay over 96 hours. Overexpression of CCR2 alone did not affect ONS76 cell proliferation. However, upon stimulation with 50ng/mL of CCL2, ONS76 OE-CCR2 cells exhibited a significant increase in proliferation compared to control cells and non-stimulated ONS76 OE-CCR2 (p<0.01 for all time points 24–96h). (**B**) Migration of ONS76 OE-CCR2 cells assessed by Boyden chamber assay at 16 hours. CCL2 stimulation significantly increased the migratory capacity of OE-CCR2 cells compared to unstimulated OE-CCR2 cells (p<0.0001). (**C-E**) Analysis of ERK and AKT signaling pathways in ONS76 OE-CCR2 following stimulation with 50 ng/mL of recombinant CCL2. (**C**) Representative immunoblots showing levels of phosphorylated AKT (p-AKT), total AKT, phosphorylated ERK (p-ERK), total ERK, and Actin (loading control) at various time intervals (1, 5, 15, 30, and 60 minutes) post-stimulation compared to non-stimulated (NS) cells. (**D**) Densitometry analysis of p-ERK relative to total ERK levels across three independent experiments. Stimulation with CCL2 revealed a visible tendency towards increased ERK activation, particularly at the 1-minute and 5-minute time points. (**E**) Densitometry analysis of p-AKT relative to total AKT levels. In contrast to ERK, no consistent trend in activation of the AKT pathway was observed following CCL2 treatment in these cells. Data represents results from three independent experiments and is expressed as median with interquartile range or as fold change in arbitrary units (AU). Statistical significance was determined using the non-parametric Mann-Whitney test. Significance levels: **p<0.01, ****p<0.0001.

### *CCR2* overexpression increases MB invasion and reduces survival *in vivo*

To understand the role of CCR2 in MB growth and progression *in vivo*, we orthotopically injected ONS76 cells into the cerebellum of NSG mice and assessed survival, tumor invasion and dissemination (**Figure 3A**).

**Figure 3.**
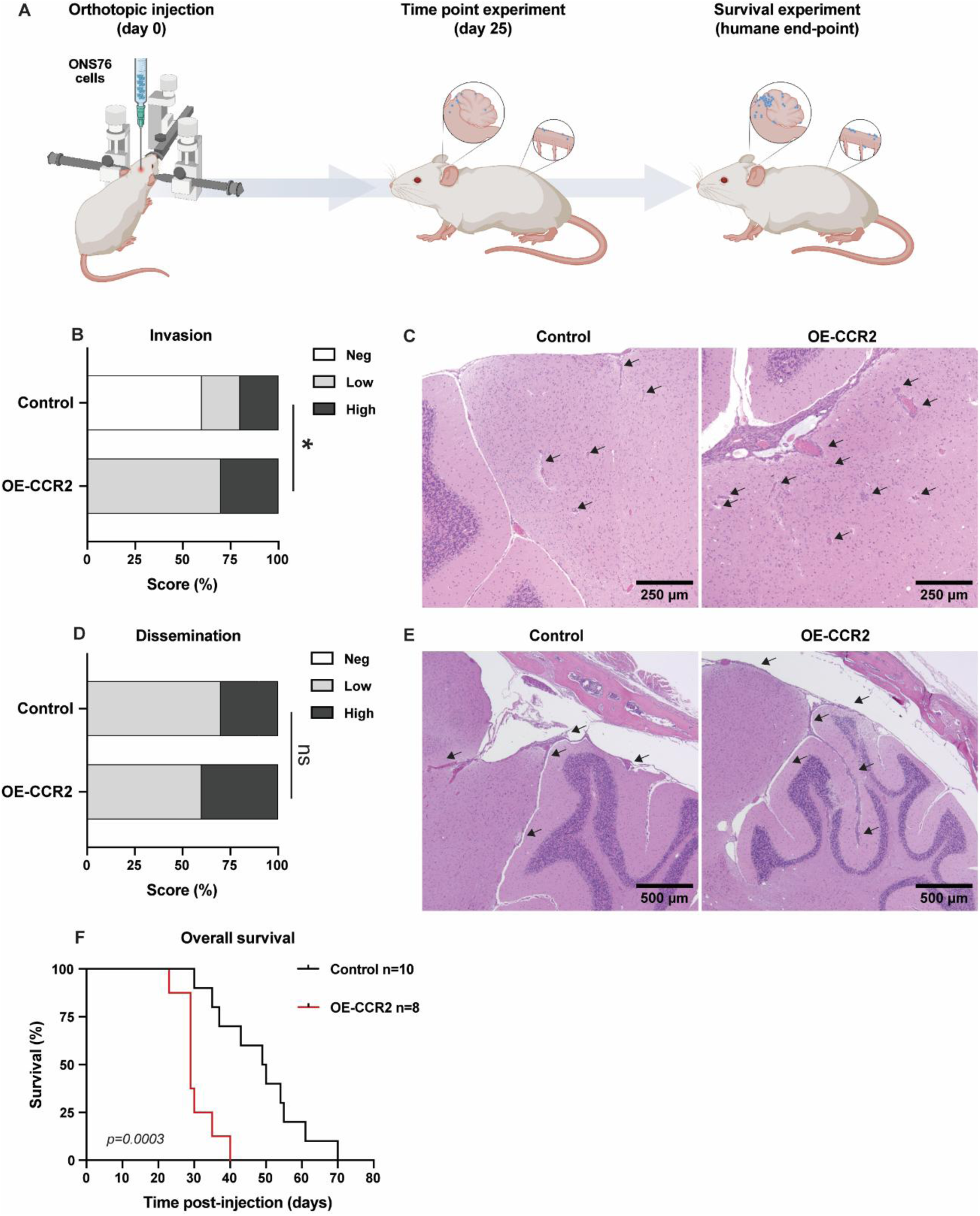
Overexpression of *CCR2* induced a more aggressive disease phenotype in orthotopic mice models. (**A**) Schematic representation of the orthotopic injection of ONS76 cells in NSG mice and experimental outline. (**B**) At 25 days post injection semi-quantitative scoring revealed higher invasion of the brain in mice injected with ONS76 OE-CCR2 cells compared to the control group. Histopathological scoring of invasion: Neg – no invasion; Low – mild invasion; High – marked brain invasion by cancer cells. (**C**) Representative images of brain invasion in both groups of mice. Arrows depict invasion *foci*. Scale bar: 250μm. (**D**) Semi-quantitative scoring did not reveal differences in dissemination between the two groups. Histopathological scoring of dissemination: Neg – no dissemination; Low – mild dissemination; High – marked dissemination in the leptomeninges by cancer cells. (**E**) Representative images of brain dissemination in both groups. Arrows depict dissemination areas. Scale bar: 500μm (**F**) Survival analysis revealed that mice injected with ONS76 OE-CCR2 cells (n=8) have decreased survival when compared to control mice (n=10) (p=0.0003). Data is expressed as percentage. Statistical significance was determined using the Ordinal Logistics Regression and Log-rank Mantel-Cox tests. Significance levels: *p<0.05, ns – not significant.

On day 25 post-injection, histopathological analysis revealed that mice injected with ONS76 OE-CCR2 had significantly increased brain invasion compared to the control group (**Figure 3B and 3C**) and a tendency to increased leptomeningeal dissemination in the brain (**Figure 3D and 3E**) and spinal cord (**Figure S4A and S4B**). Additionally, mice with CCR2-driven tumors showed significantly reduced survival compared to the control group (*p*=0.0003; **Figure 3F**).

Collectively, these results indicate that increased *CCR2* expression associates with disease progression and reduced animal survival, consistent with the results from MB patients’ datasets.

### CCR2 inhibition with Cenicriviroc decreases cancer cell proliferation, migration and signaling *in vitro*

Given the limited therapeutic options available for Group 3 MB, and our findings identifying CCR2 as a potential therapeutic target, we performed an *in-silico* drug screening using the Drug–Target Relationship Predictor (DEcRyPT) to identify compounds targeting this receptor. Among the 10,000 compounds identified, we selected the top 50 based on their high affinity for CCR2, the ability to cross the blood-brain barrier (BBB) and their prior use in clinical trials. This list of compounds was then further narrowed down based on a literature review, specific biochemical and pharmacological properties (**Figure 4A**). We selected 3 candidate compounds for *in vitro* validation: Cenicriviroc (CVC; dual CCR2/CCR5 inhibitor), Marizomib (MAR; proteasome inhibitor) and Romidepsin (ROM; HDAC inhibitor).

**Figure 4.**
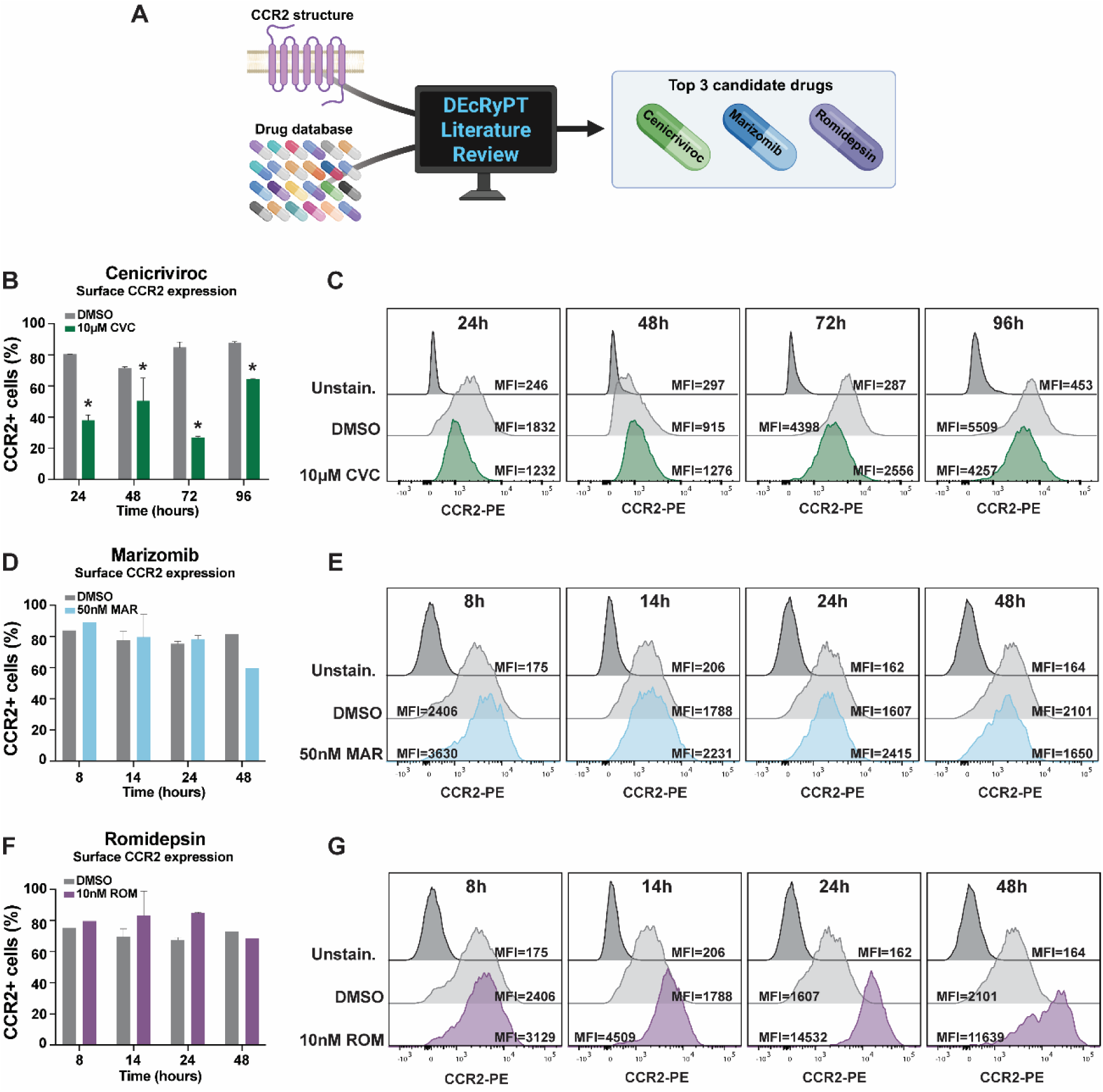
Cenicriviroc reduces CCR2 expression in MB cells. (**A**) Schematic representation of the *in-silico* drug screening approach performed to identify drugs specifically targeting CCR2. Flow cytometric evaluation of CCR2 expression in ONS76 OE-CCR2 MB cells treated with 10 µM of CVC from 24h to 96h (**B**), and representative histograms of the median fluorescence intensity (MFI) for cells treated with CVC (**C**); 50 nM of MAR from 8h to 48 hours (**D**), and representative histograms of the MFI for cells treated with MAR (**E**); and 10 nM of Romidepsin from 8h to 48 hours (**F**) and representative histograms of the MFI for cells treated with ROM (**G**). DMSO was used as a negative control across all conditions. Results reflect data from two independent experiments performed in duplicate. Quantitative data are expressed as the median with interquartile range. Statistical significance was assessed using the non-parametric Mann-Whitney test. Significance levels: *p<0.05.

First, we evaluated the ability of these compounds to decrease the levels of CCR2 at the cell surface of MB cancer cells by flow cytometry. We observed that CVC significantly reduced both the percentage of CCR2-positive cells (**Figure 4B**) and the surface expression levels of CCR2 in ONS76 (**Figure 4C**) and UW-228 OE-CCR2 cells (**Figure S5A and S5B**). MAR did not impact the percentage of CCR2-positive cells (**Figure 4D**) nor their expression levels (**Figure 4E**). In contrast, ROM increased both the percentage of CCR2-positive cells (**Figure 4F**) and CCR2 expression levels at cell surface immediately after 8h of incubation (**Figure 4G**), although without significance.

We next assessed the impact of selected compounds in cancer cell functions and in CCR2 downstream signaling. CVC effectively reduced proliferation (**Figure 5A**) and migration in ONS76 OE-CCR2 (*p*=0.0293; **Figure 5B**). Similar results were observed in UW288 OE-CCR2 cells (**Figure S5C** and **Figure S5D**). Activation of ERK in response to CCR2 stimulation with CCL2 was decreased following incubation with CVC (**Figure 5C and 5D**). In contrast, no inhibitory effect was observed on AKT activation (**Figure 5C and 5E**).

**Figure 5.**
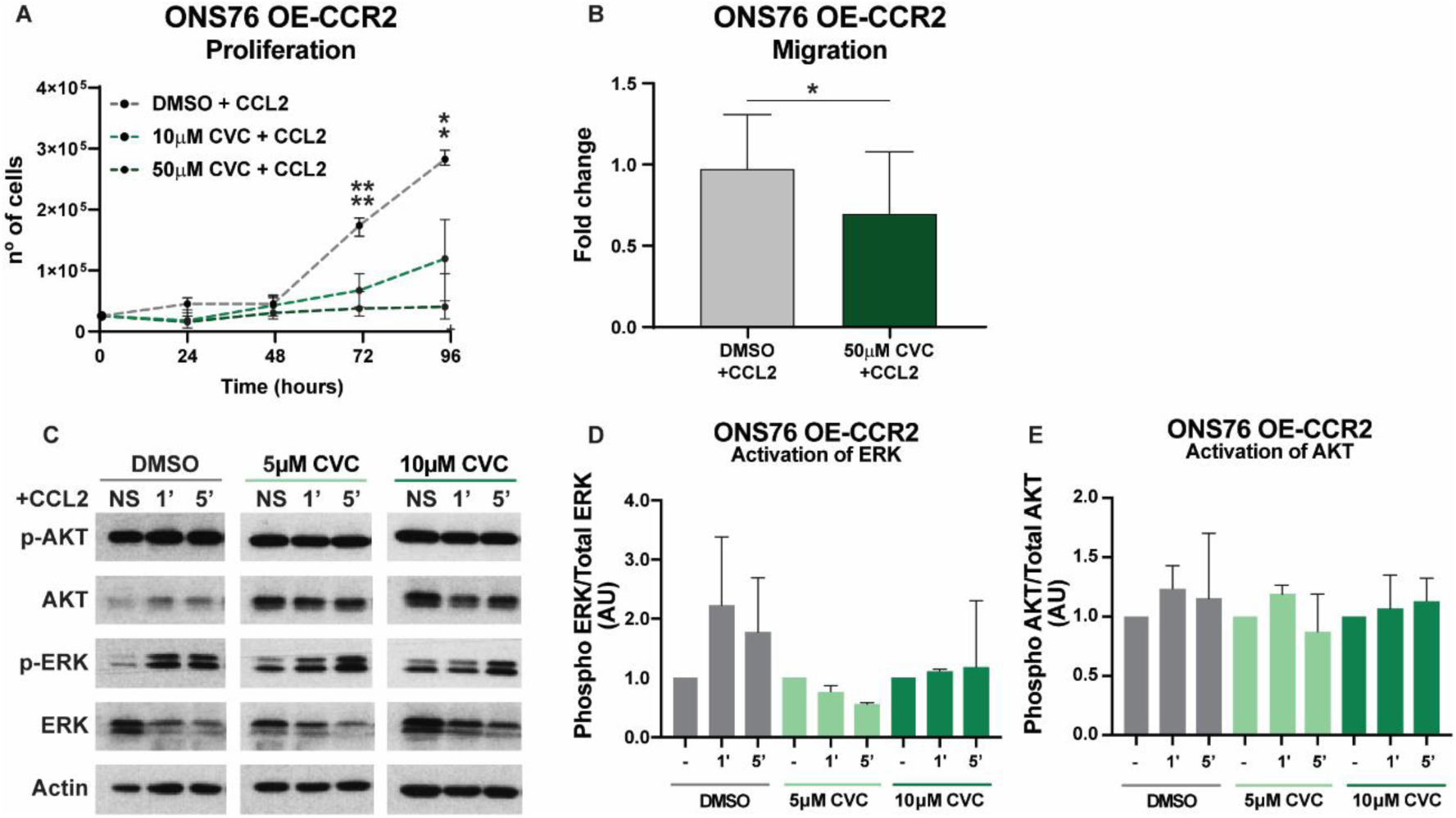
Cenicriviroc inhibits proliferation, migration and CCR2 downstream signaling in MB cells. (**A**) Dose dependent inhibition of proliferation in ONS76 OE-CCR2 cells stimulated with CCL2, assessed by MTS assay. (**B**) Incubation with 50 μM of CVC significantly reduced migration in ONS76 OE-CCR2 cells compared to the DMSO control group (p=0.0293). (**C**) Representative immunoblots for phosphorylated and total AKT/ERK after treatment with 5 μM or 10 μM CVC under CCL2 stimulation. (**D-E**) Densitometry analysis showed that CVC inhibits the ERK pathway activation in response to CCL2, but has no effect on AKT signaling. Data represents the results of three independent experiments. Cell counts and densitometry are expressed as the median with interquartile range or fold change. Statistical significance was determined using the non-parametric Mann-Whitney test. Significance levels: *p<0.05, **p<0.01.

MAR reduced proliferation of ONS76 OE-CCR2 cells after 96h of treatment (**Figure S6A**) and suppressed downstream signaling by inhibiting the activation of both ERK (**Figure S6B and S6C**) and AKT (**Figure S6B and S6D**).

Lastly, ROM significantly reduced ONS76 OE-CCR2 cell proliferation after 48h of incubation (**Figure S6E**) but did not affect ERK or AKT activation (**Figure S6F-S6H**).

Of note, the concentrations of CVC, MAR and ROM used in the *in vitro* assays did not affect the viability of ONS76 OE-CCR2 MB cells (**Figure S7A-S7C**). In UW228 MB cells, CVC significantly reduced viability after 72 and 96h of treatment (**Figure S7D**). Nevertheless, migration assays and other functional analyses were also performed at earlier time points, supporting the notion that the observed effects are unlikely to be attributable to reduced cell viability in the UW228 MB cell line.

Our *in vitro* results suggest that CVC is the most specific and effective compound to treat CCR2-driven MB.

### Cenicriviroc inhibits MB tumor development *in vivo*

Based on our *in vitro* data, CVC was subsequently evaluated *in vivo* as the most promising compound. We investigated its efficacy in treating MB using orthotopic mice models. Four days after intracranial injection of ONS76 OE-CCR2 cells in the cerebellum, NSG mice were randomized and treated by intraperitoneal injection with either vehicle (n=15) or 30mg/kg of CVC (n=10). Treatment was administered in a 5 days on/2 days off scheme for 3 weeks (**Figure 6A**). Animals were monitored every other day and no significant differences in body weight were observed between groups during treatment **(Figure 6B**). At the experimental endpoint (24h after the final treatment administration), bioluminescence imaging was performed prior to euthanasia. Treatment with CVC significantly decreased tumor burden compared to the vehicle-treated group (*p*=0.0315; **Figure 6C and 6D**). Histopathological analysis revealed a trend towards less invasion (**Figure 6E and 6F**) and dissemination of cancer cells in CVC-treated mice when compared to vehicle-treated animals (**Figure 6G and 6H**). These results show that CVC is an effective treatment for CCR2-driven MB.

**Figure 6.**
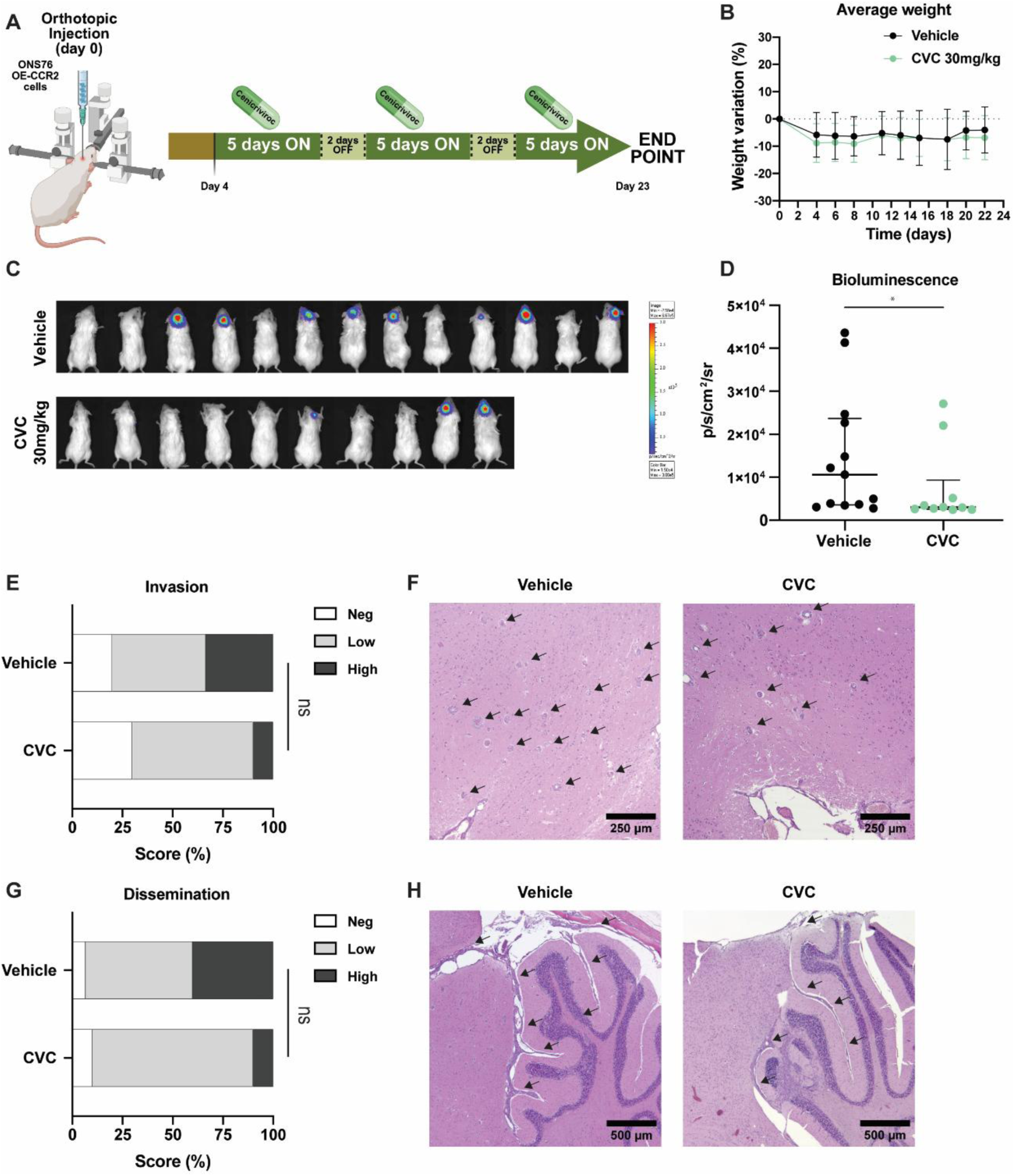
Cenicriviroc effectively reduces CCR2-driven tumor growth in an orthotopic MB mouse model. (**A**) Schematic representation of the experimental design. Mice were injected in the cerebellum with 5 × 10⁵ ONS76 OE-CCR2 cells per animal. Beginning on day 4, mice were treated with vehicle (n=15) or 30 mg/kg CVC (n=10) in cycles of 5 days ON and 2 days OFF (**B**) Average body weight variation (%) monitored every other day throughout the treatment period. (**C**) Representative whole-body bioluminescence images captured at day 24 post-injection. (**D**) Quantification of the average radiance signal (p/s/cm^2^/sr) confirmed a significant reduction in overall tumor burden in CVC-treated mice compared to vehicle controls (p=0.0315). (**E**) Semi-quantitative histopathological scoring of invasion in the brain parenchyma in CVC-treated animals. (**F**) Representative H&E-stained images of tumor invasion in both groups. Black arrows denote individual invasion foci. Scale bar: 250 μm. (**G**) Semi-quantitative analysis of tumor dissemination into the leptomeningeal compartment in mice treated with CVC relative to the vehicle cohort. (**H**) Representative images depicting areas of brain dissemination (indicated by arrows). Scale bar: 500 μm. Quantitative data in panel D are expressed as individual data points with the median and interquartile range, while scoring data in panels E and G are expressed as percentages of the respective cohorts. Statistical significance was determined using the non-parametric Mann-Whitney and the Ordinal Logistics Regression tests. Significance levels: *p<0.05.

## Discussion

Effective therapies for Group 3 MB remain scarce, highlighting a critical need for improved prognostic tools and therapeutic approaches. Identifying novel biomarkers is a key step towards a more precise patient stratification and personalized therapy. Moreover, developing therapeutic strategies targeting both primary and disseminated disease is crucial to improve patient survival as current options often fail to address the complexity and aggressiveness of metastatic disease.

Recent studies have associated the CCL2/CCR2 pathway with MB dissemination, with supporting evidence emerging from animal models and clinical patient cohorts [18, 19, 22]. While high *CCL2* expression has been linked to decreased survival in recurrent MB patients [17], the impact of *CCR2* expression in MB patient outcomes had not previously been explored. In this study, we report an association between high *CCR2* expression and decreased survival among pediatric patients with metastatic Group 3 MB, particularly the γ subtype, known to have the most unfavorable prognosis [35]. Furthermore, analysis of an independent cohort of MB patients containing paired primary and metastatic or recurrent tumor samples, showed that CCR2 levels are increased in most metastatic and recurrent tumors. Collectively, these observations highlight the role of CCR2 in MB progression and support its utility as both a prognostic biomarker and a candidate therapeutic target in high-risk MB molecular subtypes.

Activation of CCR2 by its primary ligand CCL2 leads to rapid changes in the cell phosphoproteome through the activation of the MAPK/ERK and PI3K/AKT signaling pathways [39–42]. These pathways are relevant in MB, as they are implicated in tumor growth, stemness, therapy resistance, and as targets for combination therapies [43, 44]. CCL2/CCR2 signaling has been shown to promote tumor cell proliferation and migration in various malignancies [42, 45]. Consistent with the established role of this pathway in systemic malignancies, we observed that CCL2/CCR2 signaling enhances both cell proliferation and migration in MB cells, with effects that are substantially potentiated by exogenous CCL2 stimulation.

To further investigate CCR2 downstream signaling in MB, we assessed the phosphorylation levels of ERK and AKT. Consistent with the findings from Huang *et al.* (2019), who observed a peak in ERK phosphorylation 1.5 minutes after CCL2 stimulation in CCR2-engineered HEK-293T cells, we detected increased ERK phosphorylation as early as 1-minute post-CCL2 stimulation [41]. However, in contrast to Li *et al*. (2012), who reported increased AKT activation alongside MEK/ERK activation after CCL2 stimulation [42], we observed no changes in AKT phosphorylation.

Using an orthotopic mouse model of Group 3 MB, we demonstrated that CCR2 overexpression leads to increased cell invasion and extended brain and spinal cord dissemination, which ultimately reduce mice survival (ONS76 OE-CCR2 - 29 days *vs* ONS76 Control 49.5 days). These *in vivo* findings corroborate our *in vitro* observations demonstrating CCR2-mediated enhancement of MB cell proliferation and migration. Importantly, the experimental data from ONS76 xenografts aligns with clinical observations from the Cavalli *et al.* (2017) dataset [30], where elevated *CCR2* expression was detected in pediatric metastatic Group 3 MB cohort and correlated with decreased patient survival. This concordance between preclinical models and patient outcomes validates the clinical relevance of CCR2 as a driver of Group 3 MB and supports its potential as a prognostic marker and a therapeutical target.

While standard MB treatment results in excellent outcomes for WNT subgroup patients (>95% 5-year survival), Group 3 MB patients face significantly poorer prognoses (50-60% survival) [7]. Furthermore, Group 3 MB exhibits the most aggressive phenotype and is highly propense to dissemination [3, 4, 7, 30, 46]. Therapy is further challenging given the genomic divergence between metastatic and primary tumors [47], which renders standard therapies less effective in disseminated disease. Therefore, targeting the CCL2/CCR2 pathway represents a promising approach. We performed an *in-silico* drug screening prioritizing compounds with high predicted affinity against CCR2, BBB permeability, and prior usage in the clinical setting. From this analysis, we identified 3 top candidates, CVC, MAR, and ROM. *In vitro* characterization of the impact in CCR2-overexpressing MB cells revealed distinct mechanisms of action among these compounds. CVC, a dual CCR2/CCR5 antagonist previously tested in clinical trials for non-alcoholic steatosis [48] and HIV infection [49], decreased CCR2 surface expression by 50% and inhibited cell proliferation and migration. In ONS76 OE-CCR2 cells, CVC treatment reduced the phosphorylation of CCR2 downstream effector ERK, indicating the suppression of pro-survival and pro-migratory signaling pathways. In contrast, MAR (a proteasome inhibitor) and ROM (an HDAC inhibitor) demonstrated anti-proliferative effects that appeared independent of CCR2 inhibition at the tested concentrations. MAR did not affect CCR2 expression levels while ROM paradoxically increased CCR2 expression. This CCR2 upregulation likely reflects epigenetic remodeling induced by HDAC inhibition, which has been shown to modulate chemokine receptor expression in other contexts [50]. These findings highlight CVC as the CCR2 lead candidate antagonist and therefore, it was selected for further *in vivo* validation.

In a orthotopic mouse model representative of Group 3 MB, CVC demonstrated efficacy by reducing tumor growth as well as brain invasion and dissemination. Importantly, CVC was well-tolerated, with no impact on animal weight. These findings demonstrate that pharmacological inhibition of CCR2 can effectively halt MB progression, thus supporting the clinical translation of CCR2-targeted therapies for high-risk MB.

Although our orthotopic mice models effectively replicate MB dissemination, they lack a functional immune system. Since CCL2/CCR2 pathway regulates immune cell recruitment, and CCR2 inhibition has been shown to synergize with immunotherapies in other cancer types [28, 29], evaluating this therapeutic strategy in immunocompetent models may reveal additional therapeutic benefits. Our findings demonstrate that increased CCR2 expression is associated with poorer survival outcomes in pediatric patients with Group 3 MB and promotes a more aggressive phenotype in preclinical models of the disease. These results highlight CCR2 as a prognostic biomarker and a potential therapeutic target for high-risk MB. Notably, inhibition of CCR2 by CVC effectively reduced tumor growth, invasion, and dissemination *in vivo*. Importantly, prior inclusion of CVC in clinical trials for other indications [48, 49, 51] facilitates its repurposing and clinical translation for MB. Together, our data and the unmet medical need for novel Group 3 MB therapies strongly support the clinical evaluation of CCR2 inhibition, either as monotherapy or in combination with standard of care treatments, to improve the outcomes of children with this devastating disease.

Overall, our findings establish CCR2 as both a clinically relevant prognostic biomarker and an actionable target for Group 3 MB therapy. Here, we link increased CCR2 expression to metastatic progression and decreased patient survival in Group 3 tumors. This association is further corroborated by the phenotypes observed in our orthotopic mouse models. Finally, we demonstrate that CVC can be repurposed towards MB treatment by showing that CCR2 inhibition with CVC can halt tumor progression *in vivo*, thus providing a rationale for including CCR2-targeted therapy as a novel alternative for the treatment of this high-risk subgroup of patients.

## Acknowledgments

The authors would like to acknowledge GIMM scientific platforms, including the Bioimaging Platform, funded by PPBI-POCI-01-0145-FEDER-022122, the Flow Cytometry Platform, the Histopathology Platform and the Biobank, at the Centro Académico de Medicina de Lisboa (Biobanco-GIMM CAML), for their technical support. Additionally, we would like to acknowledge the Rodent Platform for animal care and technical support. Finally, the authors acknowledge the patients who kindly provided the tumor specimens used to perform immunohistochemistry.

C.C. was supported by a fellowship from Fundação para a Ciência e a Tecnologia (FCT; SFRH/BD/140299/2018 and COVID/BD/153277/2023, identified by the https://doi.org/10.54499/COVID/BD/153277/2023); R.C. was supported by an individual employment contract from FCT (ref: 2023.09105.CEECIND/CP2857/CT0006; DOI: https://doi.org/10.54499/2023.09105.CEECIND/CP2857/CT0006); and E.P. was supported by fellowships from FCT (PD/BD/128288/2017; COVID/BD/152108/2021). This study was supported by national funds through FCT, under the projects PTDC/MED-ONC/32222/2017 and 2022.08774.PTDC, Bolsa Manuel Melo, LPCC/Lions fellowship, Fundação Millennium BCP, and private donations.

This work was conducted at GIMM-CARE. GIMM-CARE is funded by the European Union under grant agreement No. 101060102. GIMM-CARE is co-funded by the Portuguese Government, FCT, ARICA – Investimentos, Participações e Gestão, Jerónimo Martins, the Gulbenkian Institute for Molecular Medicine and CAML - Lisbon Academic Medical Centre. The founders had no role in study design, data collection and analysis, decision to publish, or preparation of the manuscript.

## Authors Contribution

Conceptualization: RC and CCF. Experimental design: CC, RC, JTB and CCF. Study conduct: CC and EP. Data collection: CC, EP, IL, SN, LA and TR. Data analysis: CC, DP, PR and TR. Data interpretation: CC, RC, DP, PR, IL, RR, MR, JTB and CCF. Drafting manuscript: CC, RC and CCF. Revising and editing manuscript content: CC, EP, RC, DP, PR, TR, IL, RR, SN, NQ, LA, JM, MR, JTB and CCF. All authors approved the final version of this manuscript and take responsibility for the integrity of the data analysis and interpretation of results.

---

**Figure S1.**
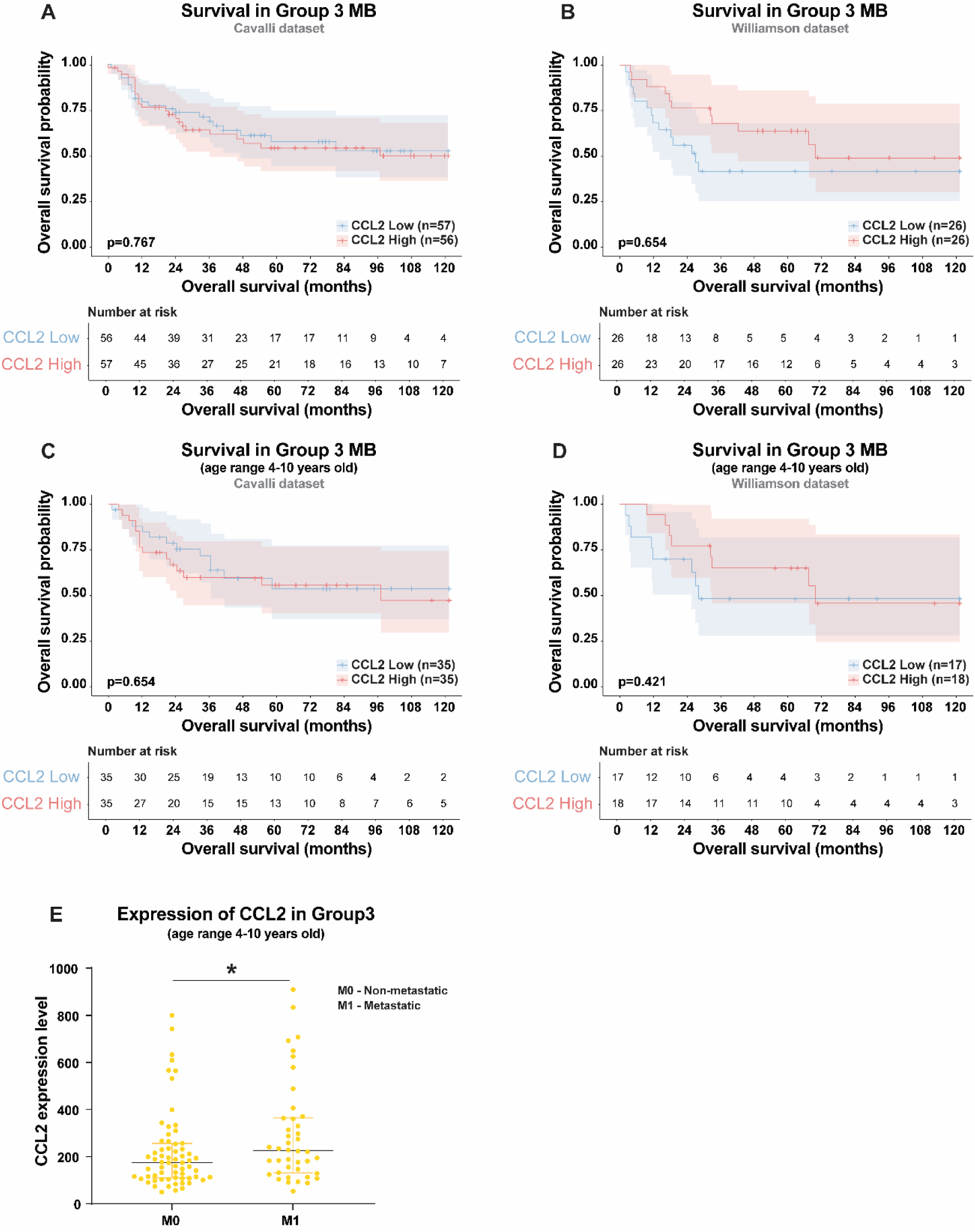
Clinical outcomes stratified by CCL2 expression levels. Kaplan-Meier analysis of overall survival (OS) stratified by CCL2 expression levels across Group 3 MB at all ages in the Cavalli **(A)** and Williamson **(B)** datasets. Kaplan-Meier analysis of OS, stratified by CCL2 expression levels, across Group 3 MB int the 4 - 10 years old age range in the Cavalli **(C)** and Williamson **(D)** datasets. Shaded areas represent 95% confidence intervals and statistical significance was determined using the two-tailed log-rank test. (**E**) CCL2 showed higher expression in metastatic (M1) patients when compared to non-metastatic (M0) ones (p=0.0361). Statistical significance was determined using the two-tailed log-rank test. Shaded areas in the KM plots represent 95% confidence intervals. Number-at-risk tables are provided below each survival curve to illustrate the cohort size and remaining patients at each 12-month interval. In the expression analysis in panel G, individual data points are presented with the median with interquartile range. Differences were considered statistically significant for p-values <0.05, according to the non-parametric Mann-Whitney test and Log-rank Mantel-Cox test. Significance levels: *p<0.05.

**Figure S2.**
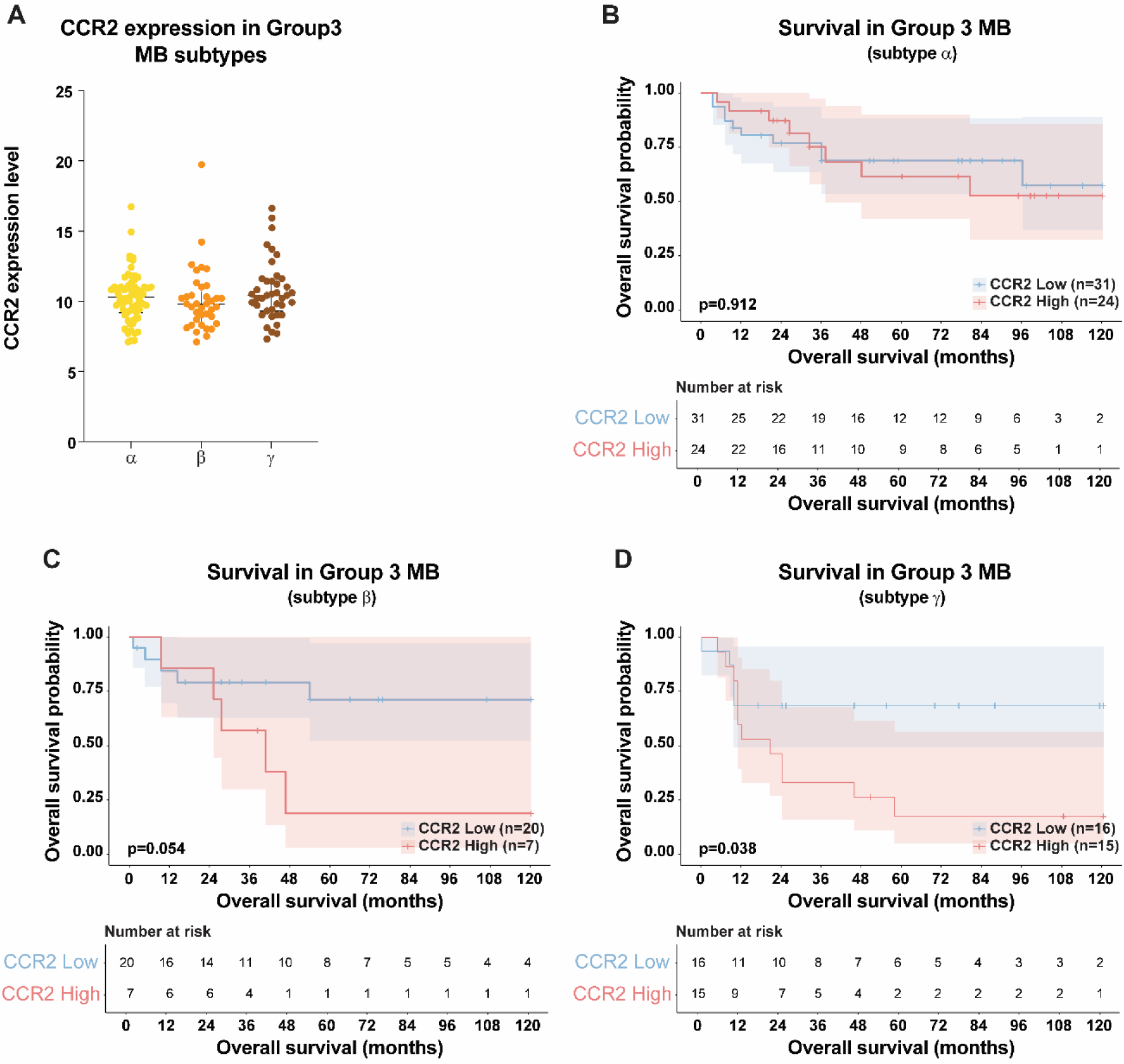
*CCR2* gene expression associates with Group 3 subtype γ worse prognosis. (**A**) *CCR2* gene expression across Group3 MB subtypes (Kruskal-Wallis p=0.19). Kaplan-Meier analysis of survival stratified according to CCR2 gene expression across Group 3 MB (**B**) subtype α, (**C**) subtype β and (**D**) subtype γ. Shaded areas in the Kaplan-Meier plots represent 95% confidence intervals, and vertical tick marks indicate censored data. Statistical analysis was performed using Kruskal-Wallis testing and Log-rank Mantel-Cox test.

**Figure S3.**
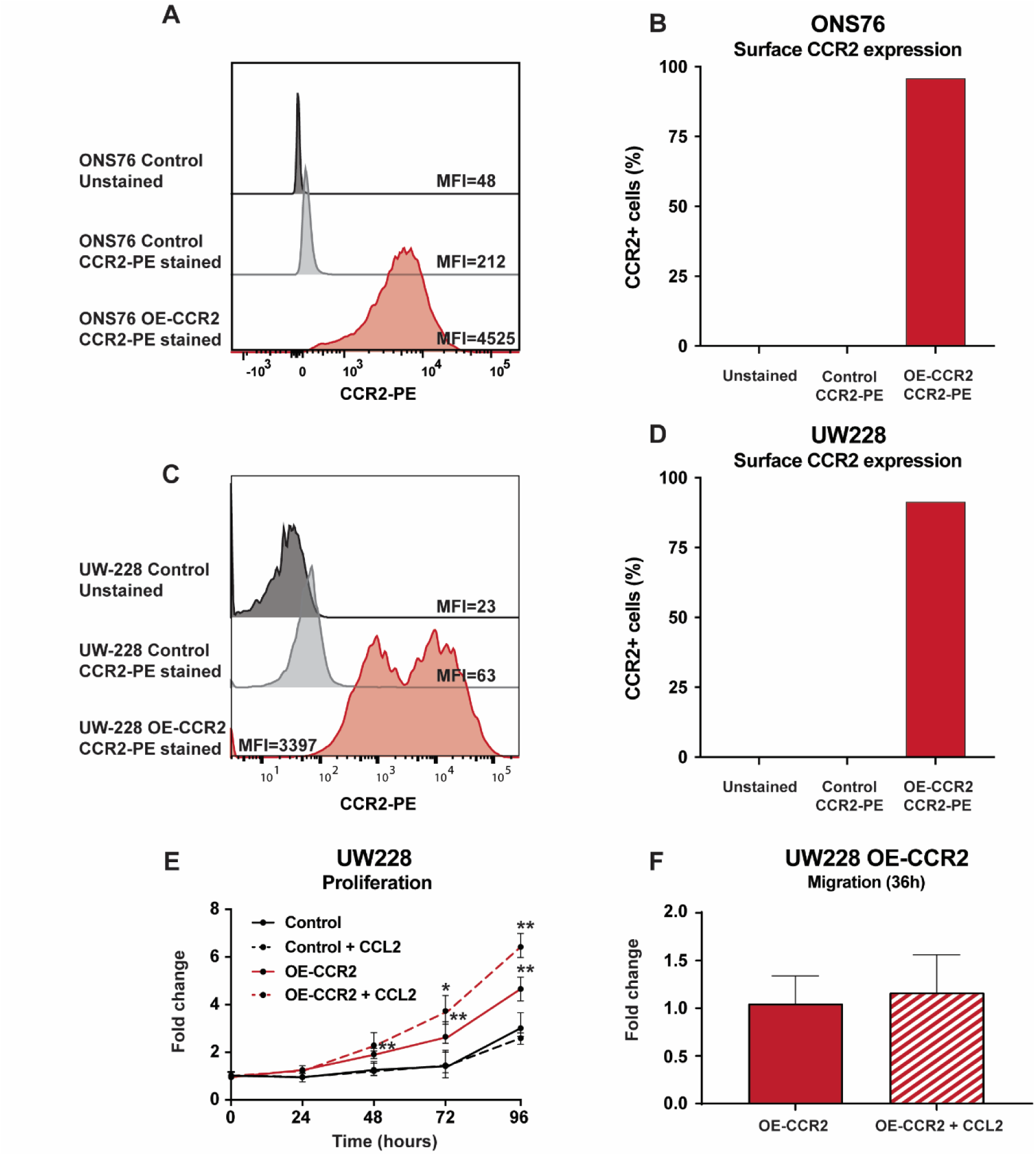
Validation of CCR2 surface expression in MB cell and functional validation of CCR2 overexpression in UW228 MB cells. (**A**) Representative flow cytometry histograms for ONS76 cells showing the distribution of CCR2-PE staining. The stable overexpression (OE-CCR2) line exhibited a marked shift in median fluorescence intensity (MFI = 4525) compared to the vector control (MFI = 212) and unstained control (MFI = 48). (**B**) Quantification of the percentage of CCR2-positive ONS76 cells, demonstrating near-total surface expression in the OE-CCR2 population compared to controls. (**C**) Representative histograms for UW228 cells showing a similar substantial shift in fluorescence intensity in OE-CCR2 cells (MFI = 3397) relative to vector control (MFI = 63) and unstained control (MFI = 23) populations. (**D**) Quantification of the percentage of CCR2-positive UW228 cells, confirming high-efficiency surface expression following stable transfection (**E**) Proliferation kinetics of UW228 cells with and without stimulation with 50 ng/mL CCL2 (p<0.05 at 72h; p<0.01 at 96h). (**F**) Migration assay in UW228 OE-CCR2 cells at 36 hours. Data represents the results of three independent experiments performed in triplicate. Values are expressed as the median with the interquartile range. Statistical significance was determined using the non-parametric Mann-Whitney test. Significance levels: *p<0.05, **p<0.01.

**Figure S4.**
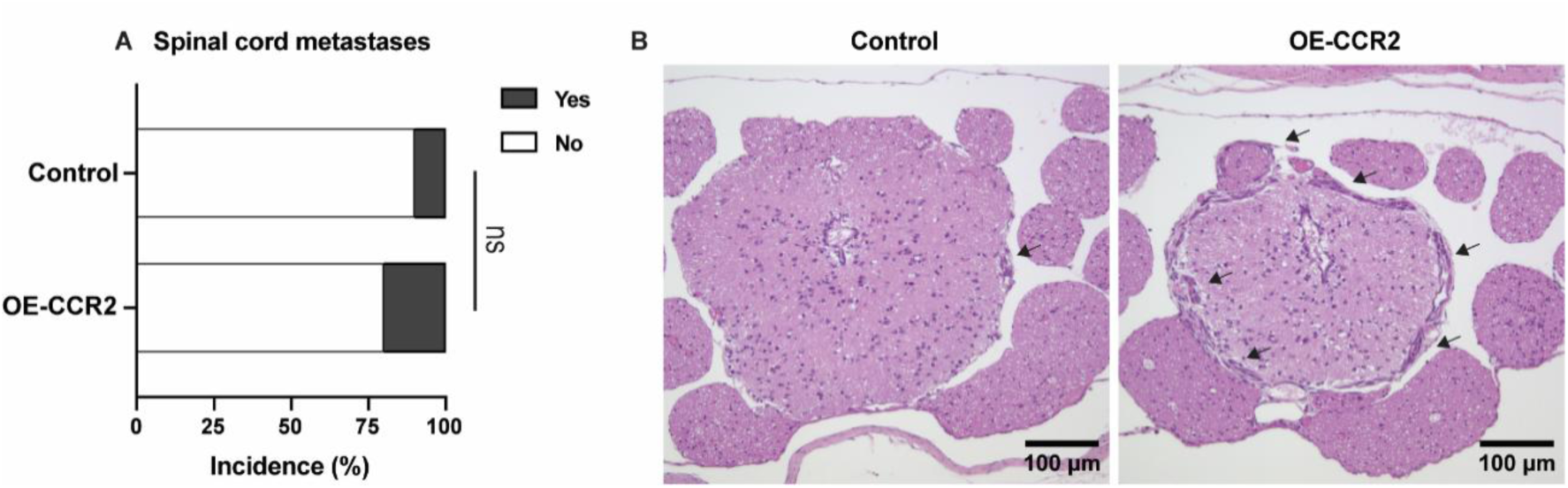
Incidence of spinal cord metastases in CCR2-driven mouse models. (**A**) Histopathological analysis showed that the incidence of spinal cord metastases is similar between ONS76 control (1 out of 10) mice and ONS76 OE-CCR2 animals (1 out of 8); **(B)** Representative images of the spinal cord of animals injected with these cells. Arrows depict metastases foci. Scale bar: 100μm. Data are expressed as percentage. Statistical significance was determined using the Ordinal Logistics Regression test. Significance levels: ns – not significant.

**Figure S5.**
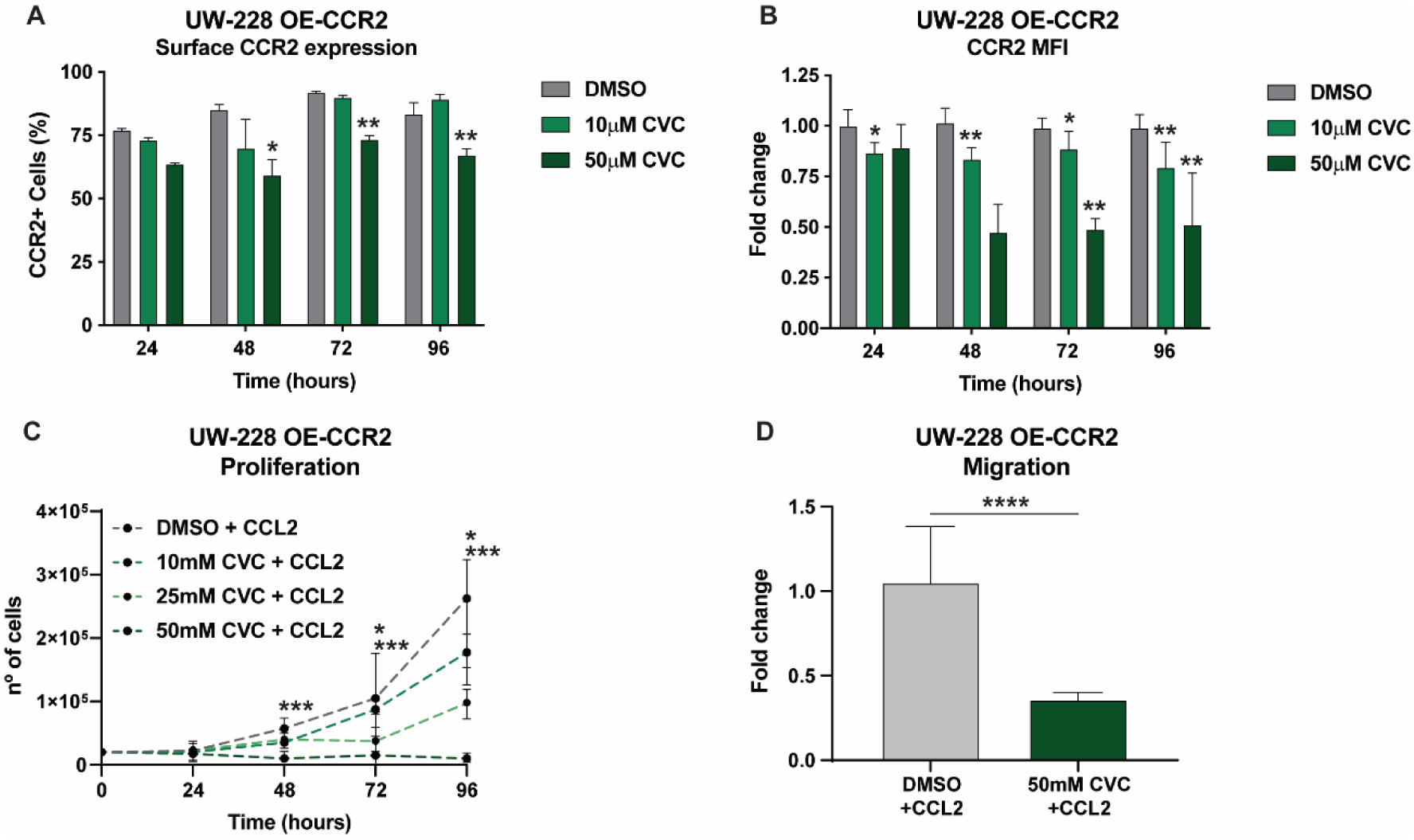
Cenicriviroc reduces CCR2 expression and inhibits proliferation and migration in UW-228 OE-CCR2 MB cells. (**A**) The percentage of CCR2 positive cells was significantly reduced after incubation with 50μM of CVC, as quantified by flow cytometry. (**B**) Expression levels of CCR2 at cell surface decreased with both 10μM and 50μM of CVC. (**C**) Dose-dependent inhibition of proliferation was observed in UW228 OE-CCR2 cells. (**D**) Effect of CVC treatment over CCL2-stimulated UW-228 OE-CCR2 MB cell migration. Results from 2 independent experiments performed in duplicates. Data are expressed as median with interquartile range. Data are expressed as median with interquartile range. Statistical significance was determined using two-tailed Mann-Whitney U tests. Significance levels: *p<0.05, **p<0.01.

**Figure S6.**
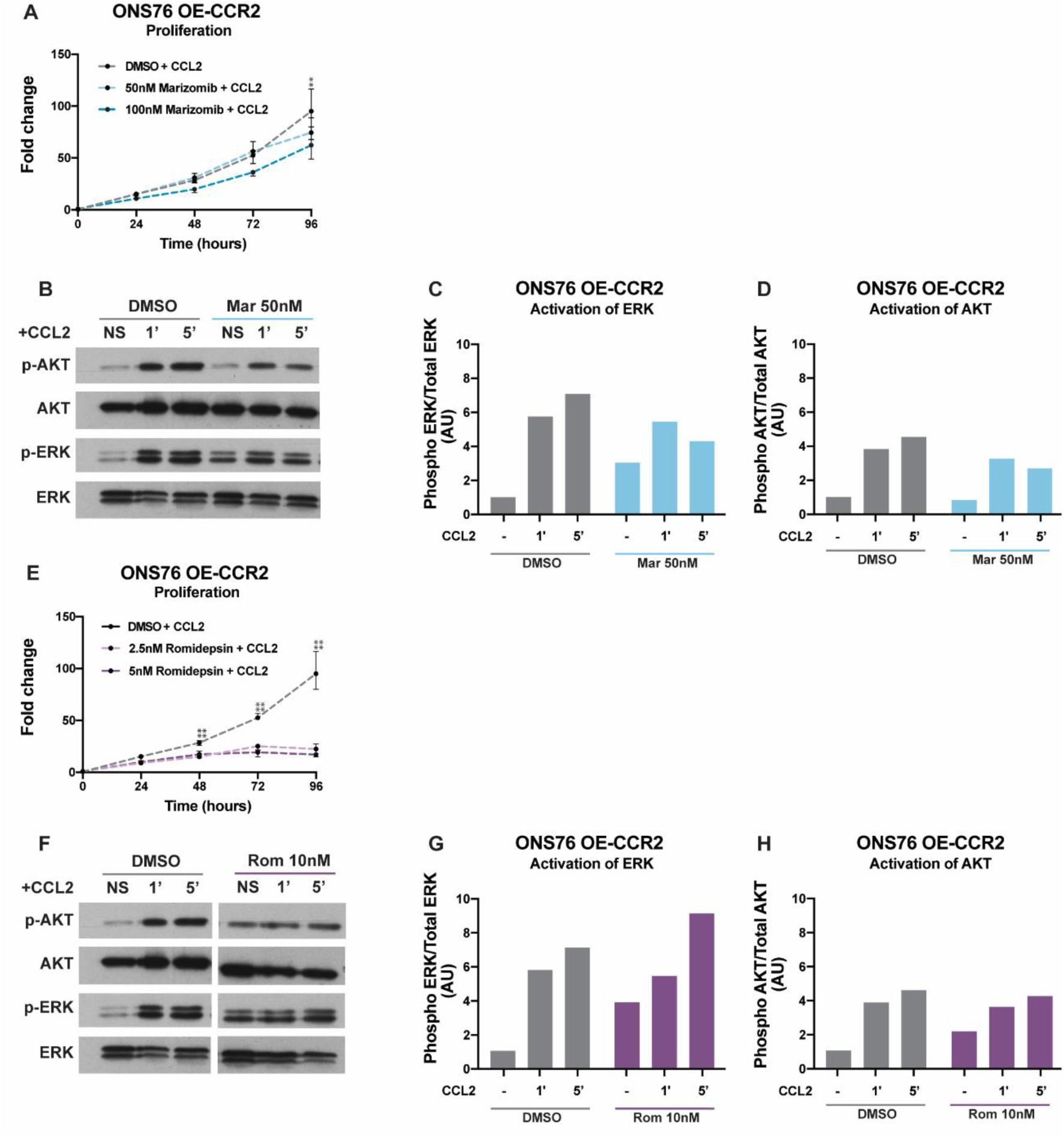
Effect of Marizomib and Romidepsin in ONS76 OE-CCR2 cell proliferation and CCR2 downstream signaling. (**A**) Incubation of ONS76 OE-CCR2 cells with 50nM and 100nM of Marizomib decreased cell proliferation at 96h, as assessed by MTS assay. (**B**) Representative immunoblots of Marizomib effect on ERK and AKT activation. Densitometry analysis revealed a reduction in ERK (**C**) and AKT (**D**) activation upon treatment of CCL2-stimulated ONS76 OE-CCR2 cells. (**E**) Both 2.5nM and 5nM of Romidepsin significantly decreased proliferation after 48h. (**F**) Representative immunoblots of Marizomib effect on ERK and AKT activation. Densitometry analysis revealed no effect on ERK (**G**) or AKT (**H**) activation upon stimulation of ONS76 OE-CCR2 with CCL2. Results from 2 independent experiments performed. Data are expressed as median with interquartile range. Statistical significance was determined using two-tailed Mann-Whitney U tests. Significance levels: *p<0.05, **p<0.01.

**Figure S7.**
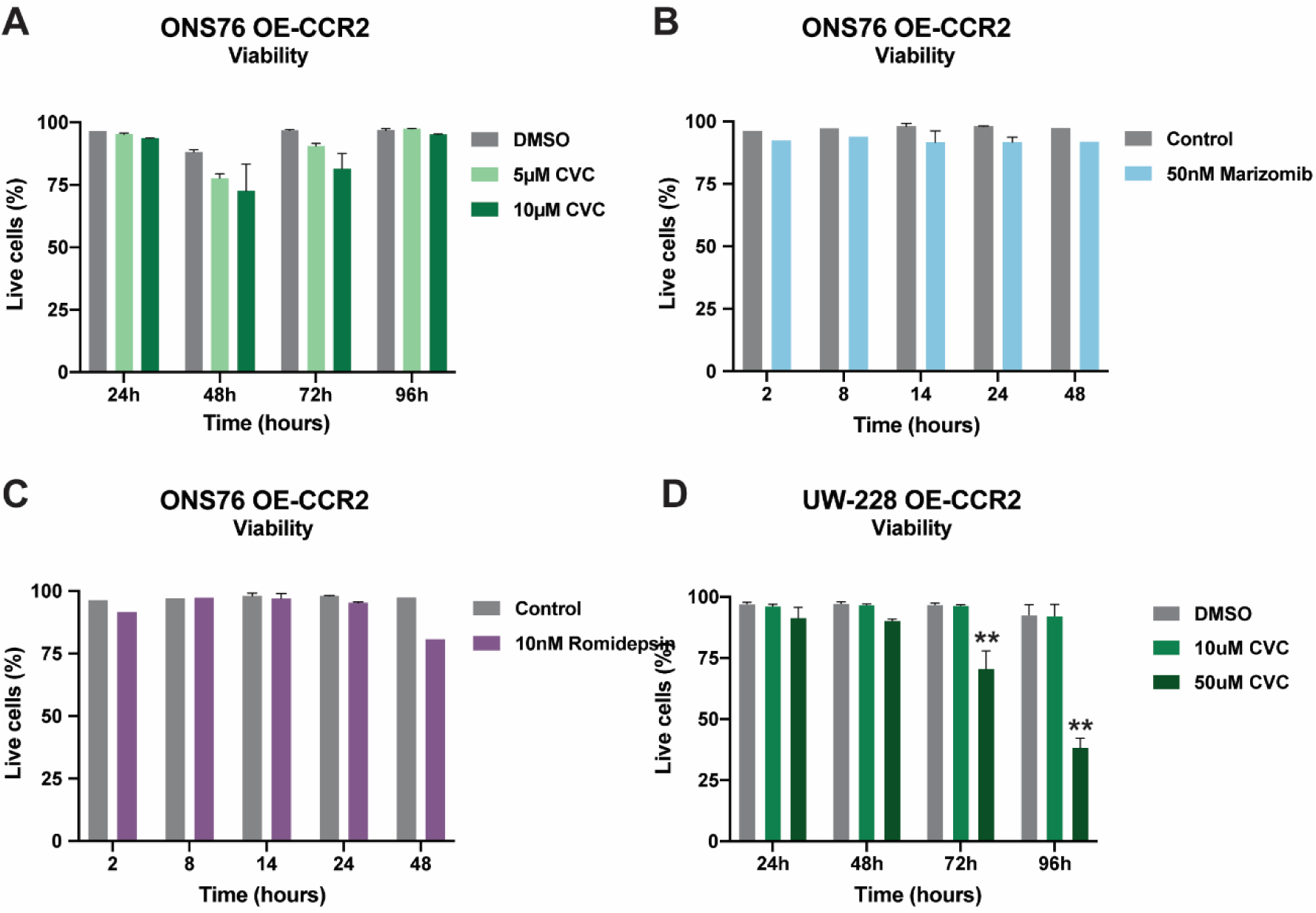
Effect of Cenicriviroc, Marizomib and Romidepsin in MB cancers viability. ONS76 OE-CCR2 cells viability was assessed by flow cytometry, after incubation with (**A**) 5 μM and 10 μM of CVC (**B**) 50nM of Marizomib or (**C**) 10nM of Romidepsin. (**D**) UW288 OE-CCR2 cells viability was assessed by flow cytometry after incubation with 10 μM and 50μM of CVC. Results from 2 independent experiments performed in triplicates. Data expressed as median with interquartile range. Data are expressed as the fold change relative to the DMSO control, shown as the median with interquartile range. All measurements were obtained via flow cytometry. Statistical significance was determined using the non-parametric Mann-Whitney test. Significance levels: *p<0.05, **p<0.01.

**Table S1.** Paired MB patient cohort description. Age intervals expressed as median with interquartile range.

| Paired MB samples collected from 1999 to 2019 (n=11) |  |  |  |
| --- | --- | --- | --- |
|  |  | N | Median age (years) |
| Sex | Male | 8 | 11.5 (6.25 - 26.5) |
|  | Female | 3 | 16 (0 - 24) |
|  | Total | 11 | 12 (6 - 24) |
| Age interval<br>(median) | 0-3 | 1 | 6 months |
|  | 4-16 | 7 | 11 (6 - 13) |
|  | 16+ | 3 | 31 (24 - 34) |
| Metastasis at diagnosis |  | 1 | 6 months |
| Type of disease | Metastatic | 3 | 12 (4 - 42) |
|  | Recurrent | 8 | 12 (6.25 - 22) |

## References

1. Mahapatra S, Amsbaugh MJ (2026) Medulloblastoma. https://www.ncbi.nlm.nih.gov/books/NBK431069/. Accessed 4 Sep 2026

2. Price M, Ballard CAP, Benedetti JR, et al (2025) CBTRUS Statistical Report: Primary Brain and Other Central Nervous System Tumors Diagnosed in the United States in 2018–2022. Neuro Oncol 27:iv1–iv66. 10.1093/neuonc/noaf194

3. Northcott PA, Korshunov A, Witt H, et al (2011) Medulloblastoma Comprises Four Distinct Molecular Variants. Journal of Clinical Oncology 29:1408–1414. 10.1200/JCO.2009.27.4324

4. Taylor MD, Northcott PA, Korshunov A, et al (2012) Molecular subgroups of medulloblastoma: the current consensus. Acta Neuropathol 123:465–472. 10.1007/s00401-011-0922-z

5. Doger de Spéville E, Kieffer V, Dufour C, et al (2021) Neuropsychological consequences of childhood medulloblastoma and possible interventions: A review. Neurochirurgie 67:90–98. 10.1016/j.neuchi.2018.03.002

6. Wang Q, Xin X, Dai Q, et al (2023) Medulloblastoma targeted therapy: From signaling pathways heterogeneity and current treatment dilemma to the recent advances in development of therapeutic strategies. Pharmacol Ther 250:108527. 10.1016/j.pharmthera.2023.108527

7. Ramaswamy V, Remke M, Bouffet E, et al (2016) Risk stratification of childhood medulloblastoma in the molecular era: the current consensus. Acta Neuropathol 131:821–31. 10.1007/s00401-016-1569-6

8. Waszak SM, Northcott PA, Buchhalter I, et al (2018) Spectrum and prevalence of genetic predisposition in medulloblastoma: a retrospective genetic study and prospective validation in a clinical trial cohort. Lancet Oncol 19:785–798. 10.1016/S1470-2045(18)30242-0

9. Cho Y-J, Tsherniak A, Tamayo P, et al (2011) Integrative genomic analysis of medulloblastoma identifies a molecular subgroup that drives poor clinical outcome. J Clin Oncol 29:1424–30. 10.1200/JCO.2010.28.5148

10. Kool M, Korshunov A, Remke M, et al (2012) Molecular subgroups of medulloblastoma: an international meta-analysis of transcriptome, genetic aberrations, and clinical data of WNT, SHH, Group 3, and Group 4 medulloblastomas. Acta Neuropathol 123:473–484. 10.1007/s00401-012-0958-8

11. Zeltzer PM, Boyett JM, Finlay JL, et al (1999) Metastasis stage, adjuvant treatment, and residual tumor are prognostic factors for medulloblastoma in children: conclusions from the Children’s Cancer Group 921 randomized phase III study. J Clin Oncol 17:832–45. 10.1200/JCO.1999.17.3.832

12. Zhou W-T, Wu T, Lu Y-F, et al (2026) Development of an integrated risk stratification model for metastatic medulloblastoma (M2/3) using clinical, radiologic, and molecular variables. Neurooncol Adv 8:vdaf265. 10.1093/noajnl/vdaf265

13. Deshmane SL, Kremlev S, Amini S, Sawaya BE (2009) Monocyte Chemoattractant Protein-1 (MCP-1): An Overview. Journal of Interferon & Cytokine Research 29:313–326. 10.1089/jir.2008.0027

14. Tsuyada A, Chow A, Wu J, et al (2012) CCL2 Mediates Cross-talk between Cancer Cells and Stromal Fibroblasts That Regulates Breast Cancer Stem Cells. Cancer Res 72:2768–2779. 10.1158/0008-5472.CAN-11-3567

15. Yoshimura T, Li C, Wang Y, Matsukawa A (2023) The chemokine monocyte chemoattractant protein-1/CCL2 is a promoter of breast cancer metastasis. Cell Mol Immunol 20:714–738. 10.1038/s41423-023-01013-0

16. Garzia L, Kijima N, Morrissy AS, et al (2018) A Hematogenous Route for Medulloblastoma Leptomeningeal Metastases. Cell 172:1050–1062.e14. 10.1016/j.cell.2018.01.038

17. Liu H, Sun Y, O’Brien JA, et al (2020) Necroptotic astrocytes contribute to maintaining stemness of disseminated medulloblastoma through CCL2 secretion. Neuro Oncol 22:625–638. 10.1093/neuonc/noz214

18. Low SYY, Bte Syed Sulaiman N, Tan EEK, et al (2020) Cerebrospinal fluid cytokines in metastatic group 3 and 4 medulloblastoma. BMC Cancer 20:554. 10.1186/s12885-020-07048-0

19. Chen Y, Zhao H, Zhang H, et al (2024) Cytokine profile of cerebrospinal fluid in pediatric patients with metastatic medulloblastoma. Heliyon 10:e38504. 10.1016/j.heliyon.2024.e38504

20. Fein MR, He X-Y, Almeida AS, et al (2020) Cancer cell CCR2 orchestrates suppression of the adaptive immune response. Journal of Experimental Medicine 217:. 10.1084/jem.20181551

21. Wang Z, Xie H, Zhou L, et al (2016) CCL2/CCR2 axis is associated with postoperative survival and recurrence of patients with non-metastatic clear-cell renal cell carcinoma. Oncotarget 7:51525–51534. 10.18632/oncotarget.10492

22. Brummer G, Fang W, Smart C, et al (2020) CCR2 signaling in breast carcinoma cells promotes tumor growth and invasion by promoting CCL2 and suppressing CD154 effects on the angiogenic and immune microenvironments. Oncogene 39:2275–2289. 10.1038/s41388-019-1141-7

23. Fang W Bin, Sofia Acevedo D, Smart C, et al (2021) Expression of CCL2/CCR2 signaling proteins in breast carcinoma cells is associated with invasive progression. Sci Rep 11:8708. 10.1038/s41598-021-88229-0

24. Yang Q, Zhang J, Zhang X, et al (2021) C-C motif chemokine ligand 2/C-C receptor 2 is associated with glioma recurrence and poor survival. Exp Ther Med 21:564. 10.3892/etm.2021.9996

25. Loberg RD, Day LL, Harwood J, et al (2006) CCL2 is a potent regulator of prostate cancer cell migration and proliferation. Neoplasia 8:578–86. 10.1593/neo.06280

26. Lindholm PF, Sivapurapu N, Jovanovic B, Kajdacsy-Balla A (2015) Monocyte-Induced Prostate Cancer Cell Invasion is Mediated by Chemokine ligand 2 and Nuclear Factor-κB Activity. J Clin Cell Immunol 6:. 10.4172/2155-9899.1000308

27. Li X, Yao W, Yuan Y, et al (2017) Targeting of tumour-infiltrating macrophages via CCL2/CCR2 signalling as a therapeutic strategy against hepatocellular carcinoma. Gut 66:157–167. 10.1136/gutjnl-2015-310514

28. Flores-Toro JA, Luo D, Gopinath A, et al (2020) CCR2 inhibition reduces tumor myeloid cells and unmasks a checkpoint inhibitor effect to slow progression of resistant murine gliomas. Proceedings of the National Academy of Sciences 117:1129–1138. 10.1073/pnas.1910856117

29. Tu MM, Abdel-Hafiz HA, Jones RT, et al (2020) Inhibition of the CCL2 receptor, CCR2, enhances tumor response to immune checkpoint therapy. Commun Biol 3:720. 10.1038/s42003-020-01441-y

30. Cavalli FMG, Remke M, Rampasek L, et al (2017) Intertumoral Heterogeneity within Medulloblastoma Subgroups. Cancer Cell 31:737–754.e6. 10.1016/j.ccell.2017.05.005

31. Williamson D, Schwalbe EC, Hicks D, et al (2022) Medulloblastoma group 3 and 4 tumors comprise a clinically and biologically significant expression continuum reflecting human cerebellar development. Cell Rep 40:111162. 10.1016/j.celrep.2022.111162

32. Rodrigues T, de Almeida BP, Barbosa-Morais NL, Bernardes GJL (2019) Dissecting celastrol with machine learning to unveil dark pharmacology. Chem Commun (Camb) 55:6369–6372. 10.1039/c9cc03116b

33. Rodrigues T, Werner M, Roth J, et al (2018) Machine intelligence decrypts β-lapachone as an allosteric 5-lipoxygenase inhibitor. Chem Sci 9:6899–6903. 10.1039/c8sc02634c

34. Okamoto M, Toyama M, Baba M (2020) The chemokine receptor antagonist cenicriviroc inhibits the replication of SARS-CoV-2 in vitro. Antiviral Res 182:104902. 10.1016/j.antiviral.2020.104902

35. Morin A, Soane C, Pierce A, et al (2020) Proteasome inhibition as a therapeutic approach in atypical teratoid/rhabdoid tumors. Neurooncol Adv 2:vdaa051. 10.1093/noajnl/vdaa051

36. Vitanza NA, Biery MC, Myers C, et al (2021) Optimal therapeutic targeting by HDAC inhibition in biopsy-derived treatment-naïve diffuse midline glioma models. Neuro Oncol 23:376–386. 10.1093/neuonc/noaa249

37. Kruger AJ, Fuchs BC, Masia R, et al (2018) Prolonged cenicriviroc therapy reduces hepatic fibrosis despite steatohepatitis in a diet-induced mouse model of nonalcoholic steatohepatitis. Hepatol Commun 2:529–545. 10.1002/hep4.1160

38. Kwiatkowski K, Ciapała K, Rojewska E, et al (2020) Comparison of the beneficial effects of RS504393, maraviroc and cenicriviroc on neuropathic pain-related symptoms in rodents: behavioral and biochemical analyses. Int Immunopharmacol 84:106540. 10.1016/j.intimp.2020.106540

39. Ko J, Yun C-Y, Lee J-S, et al (2007) p38 MAPK and ERK activation by 9-cis-retinoic acid induces chemokine receptors CCR1 and CCR2 expression in human monocytic THP-1 cells. Exp Mol Med 39:129–138. 10.1038/emm.2007.15

40. Jiménez-Sainz MC, Fast B, Mayor F, Aragay AM (2003) Signaling Pathways for Monocyte Chemoattractant Protein 1-Mediated Extracellular Signal-Regulated Kinase Activation. Mol Pharmacol 64:773–782. 10.1124/mol.64.3.773

41. Huang C, Foster SR, Shah AD, et al (2020) Phosphoproteomic characterization of the signaling network resulting from activation of the chemokine receptor CCR2. Journal of Biological Chemistry 295:6518–6531. 10.1074/jbc.RA119.012026

42. Li M-Q, Li H-P, Meng Y-H, et al (2012) Chemokine CCL2 enhances survival and invasiveness of endometrial stromal cells in an autocrine manner by activating Akt and MAPK/Erk1/2 signal pathway. Fertil Steril 97:919–929.e1. 10.1016/j.fertnstert.2011.12.049

43. Badodi S, Pomella N, Lim YM, et al (2022) Combination of BMI1 and MAPK/ERK inhibitors is effective in medulloblastoma. Neuro Oncol 24:1273–1285. 10.1093/neuonc/noac052

44. Slika H, Alimonti P, Raj D, et al (2023) The Neurodevelopmental and Molecular Landscape of Medulloblastoma Subgroups: Current Targets and the Potential for Combined Therapies. Cancers (Basel) 15:3889. 10.3390/cancers15153889

45. Küper C, Beck F-X, Neuhofer W (2016) Autocrine MCP-1/CCR2 signaling stimulates proliferation and migration of renal carcinoma cells. Oncol Lett 12:2201–2209. 10.3892/ol.2016.4875

46. Schwalbe EC, Lindsey JC, Nakjang S, et al (2017) Novel molecular subgroups for clinical classification and outcome prediction in childhood medulloblastoma: a cohort study. Lancet Oncol 18:958–971. 10.1016/S1470-2045(17)30243-7

47. Wu X, Northcott PA, Dubuc A, et al (2012) Clonal selection drives genetic divergence of metastatic medulloblastoma. Nature 482:529–33. 10.1038/nature10825

48. Anstee QM, Neuschwander-Tetri BA, Wong VW-S, et al (2020) Cenicriviroc for the treatment of liver fibrosis in adults with nonalcoholic steatohepatitis: AURORA Phase 3 study design. Contemp Clin Trials 89:105922. 10.1016/j.cct.2019.105922

49. Thompson M, Saag M, DeJesus E, et al (2016) A 48-week randomized phase 2b study evaluating cenicriviroc versus efavirenz in treatment-naive HIV-infected adults with C-C chemokine receptor type 5-tropic virus. AIDS 30:869–878. 10.1097/QAD.0000000000000988

50. Wen T, Sun G, Jiang W, et al (2023) Histone deacetylases inhibitor chidamide synergizes with humanized PD1 antibody to enhance T-cell chemokine expression and augment Ifn-γ response in NK-T cell lymphoma. EBioMedicine 87:104420. 10.1016/j.ebiom.2022.104420

51. A randomized, double-blind, multicenter study to assess the safety, tolerability, and efficacy of a combination treatment of tropifexor (LJN452) and cenicriviroc (CVC) in adult patients with nonalcoholic steatohepatitis (NASH) and liver fibrosis (TANDEM). In: EU Clinical Trials Register. https://www.clinicaltrialsregister.eu/ctr-search/trial/2017-004208-24/PT. Accessed 9 Jul 2024

